# The KRAB-Zinc Finger protein ZKSCAN3 represses enhancers via embedded retrotransposons

**DOI:** 10.1101/2025.01.30.635440

**Authors:** Daniel Moore, Eugenia Wong, Charles Arnal, Stefan Schoenfelder, Mikhail Spivakov, Simon Andrews, Maria A. Christophorou

## Abstract

Gene *cis*-regulatory sequences are increasingly recognised as containing “domesticated” transposable elements that impact their function. The KRAB Zinc Finger Protein (KZFP) family of transcription factors is typically associated with transposable element silencing through establishment of heterochromatin. Here, using acute protein depletion in embryonic stem cells, we reveal that the KZFP ZKSCAN3 represses enhancer activity through targeting enhancer-embedded retrotransposons and that ZKSCAN3-mediated repression does not rely on the induction of heterochromatin. ZKSCAN3, which exhibits strong genetic association with the neurodevelopmental disorder schizophrenia, operates during neural differentiation and is necessary for proper cell specification and expression of genes that regulate axon guidance, neuronal motility and pathfinding. These findings define ZKSCAN3 as an enhancer regulator and uncover a heterochromatin-independent KZFP. Additionally, they exemplify how a KZFP epigenetically regulates enhancers in a native setting and highlight how transposable elements and their KZFP binders have shaped gene expression networks.

## Introduction

The KRAB Zinc Finger Proteins (KZFPs) are the largest family of transcription factors (TFs) in higher vertebrates^1–4^. The majority of KZFPs target Transposable Elements (TEs), mobile genetic sequences capable of transposing from one genomic location to another^3^. The considerable expansion of the KZFP family throughout evolution is thought to have been driven, at least in part, by the need to control new waves of TE insertions into genomes^5–7^. KZFPs that successfully bound and restricted the transposition of a TE were fixed and underwent positive selection; however, this produced selection pressure for TE mutations that resulted in escape from KZFP-mediated silencing, promoting the emergence of new KZFPs, in a so-called “arms race”^5,7,8^. Currently, only a small fraction of KZFPs appear necessary for the continued silencing of TEs^9–15^ and the vast majority bind to TEs that are no longer replication-competent^3^. This observation has led to the suggestion that KZFPs do not simply target TEs for silencing but facilitate the “domestication” of TE regulatory sequences that benefit the host^3,4^.

TEs are a potent source of *cis*-regulatory sequences and can act as alternative promoters, enhancers, silencers, and boundary elements^16^. The evolution of KZFPs that can dynamically bind and regulate these sequences therefore has the potential to feed into the regulation of gene expression. Indeed, KZFPs have been shown to target domesticated TEs and impact crucial aspects of mammalian biology including human embryonic genome activation^17^, adipogenesis^18^, and human brain development^19,20^, although the underlying molecular mechanisms remain poorly understood.

KZFPs contain an N-terminal KRAB domain and a C-terminal array of DNA-binding C2H2 zinc finger domains. The KRAB domain recruits the co-repressor KRAB Associated Protein 1 (KAP1), which acts as a scaffold for the assembly of heterochromatin-inducing modifiers including the H3K9 methyltransferase SETDB1, the histone deacetylase and chromatin remodelling complex NuRD, heterochromatin protein 1 (HP1) and DNA methyltransferases^21–26^. KZFPs are therefore typically thought to mediate transcriptional repression through heterochromatin induction. There have been several reports that putative TE-derived enhancers can be regulated by the dynamic induction of heterochromatin at their loci^17,27^. Furthermore, although most KZFP-targeted TEs are enriched for H3K9me3, a fraction are enriched for enhancer-associated H3K4me1 and H3K27ac^3^, and some switch between these chromatin states in different cell types, suggesting dynamic regulation. Based on these findings, KZFP-mediated enhancer regulation is assumed to occur by selective induction of heterochromatin.

ZKSCAN proteins are a poorly understood sub-class of KZFPs that contain additional N-terminal SCAN domains and exhibit a weaker affinity with the KAP1 co-repressor when assessed by mass-spectrometry^3,28^. It has therefore been hypothesised that ZKSCAN proteins, which are generally evolutionarily older, may have evolved KAP1-independent activities not associated with the induction of heterochromatin^7^. However, the mechanism of action and biological functions of this sub-family of KZFPs remain largely uncharacterised.

ZKSCAN3 (Zinc finger with KRAB and SCAN domains 3) is ZKSCAN protein that has been genetically associated with schizophrenia (SZ) in humans. Several lines of evidence converge on a functional link between the *ZKSCAN3* gene and the aetiology of this neurodevelopmental disorder. In the largest Genome Wide Association Study (GWAS) to date^29^, the top-ranked genomic region for schizophrenia risk contains the *ZKSCAN3* locus. In a follow-up study that employed targeted-exon sequencing of genes implicated in this GWAS, the *ZKSCAN3* single nucleotide polymorphism (SNP) rs733743, which produces a missense variant (Arg3Thr), was highly associated with schizophrenia risk^30^. Additionally, a separate follow-up study of SNPs identified in the GWAS found 39 SNPs to be significantly associated with SZ-discriminating changes to regional brain grey matter volume, with 9 of these SNPs being expression quantitative trait loci (eQTLs) for *ZKSCAN3*expression in the adult brain^31^. These pieces of genetic evidence suggest a functional link between ZKSCAN3 and neurodevelopment.

In this study, using acute ZKSCAN3 protein degradation and biochemical analyses, we demonstrate that ZKSCAN3 targets SINE retrotransposon sequences embedded within enhancer sites and restricts the activity of those enhancers. Notably, we find that ZKSCAN3-mediated repression occurs independently of KAP1 recruitment and H3K9me3 induction. Rather, ZKSCAN3 restricts chromatin accessibility and dampens enrichment of H3K27ac and H3K4me1, fine-tuning gene expression in a rheostat-like manner. Using single cell transcriptomic analyses of embryonic stem cells undergoing multi-lineage differentiation, we show that loss of ZKSCAN3-mediated enhancer restriction during neural specification results in aberrant expression of endodermal and mesodermal genes. Additionally, ZKSCAN3 loss leads to mis-regulation of axon guidance receptors and downstream actin cytoskeleton components associated with neuronal motility and pathfinding.

In summary, our findings demonstrate that KZFPs can regulate enhancers and modulate gene expression independently of heterochromatin induction, and suggest a mechanistic basis for the genetic association between *ZKSCAN3* and schizophrenia risk. Additionally, this study informs how KZFPs targeting domesticated TE sequences impact the regulation of gene expression networks.

## Results

### ZKSCAN3 binds to a SINE-embedded sequence within enhancers

Analysis of publicly available bulk RNA-sequencing (RNA-seq) data of early mouse embryos^32^ revealed that *Zkscan3* is expressed at multiple stages of early mammalian embryonic development, including in the pluripotent cells of the inner cell mass at E3.5 (Figure S1). To understand the function of ZKSCAN3 during early development and differentiation we therefore genetically modified mouse Embryonic Stem Cells (mESCs) to knock in a FKBPv degron tag to the C terminus of both alleles of endogenous *Zkscan3,* tagging all curated transcripts (Figures 1A, upper panel, and Figure S2) and allowing rapid depletion and recovery of the ZKSCAN3 protein. A double HA epitope tag was included to overcome limitations of antibody availability and allow ZKSCAN3 isolation and assessment of its genomic localisation. We refer to these cells as ‘*Zkscan3-Fkbpv-HA’* mESCs. We detected clear expression of the tagged protein in four independent clonal lines by immunoblot (Figure 1A, lower panel) and were able to induce degradation of the protein through the addition of the small molecule dTAG-13^33^ (‘dTAG’) to the culture medium (Figure S3).

**Figure 1.**
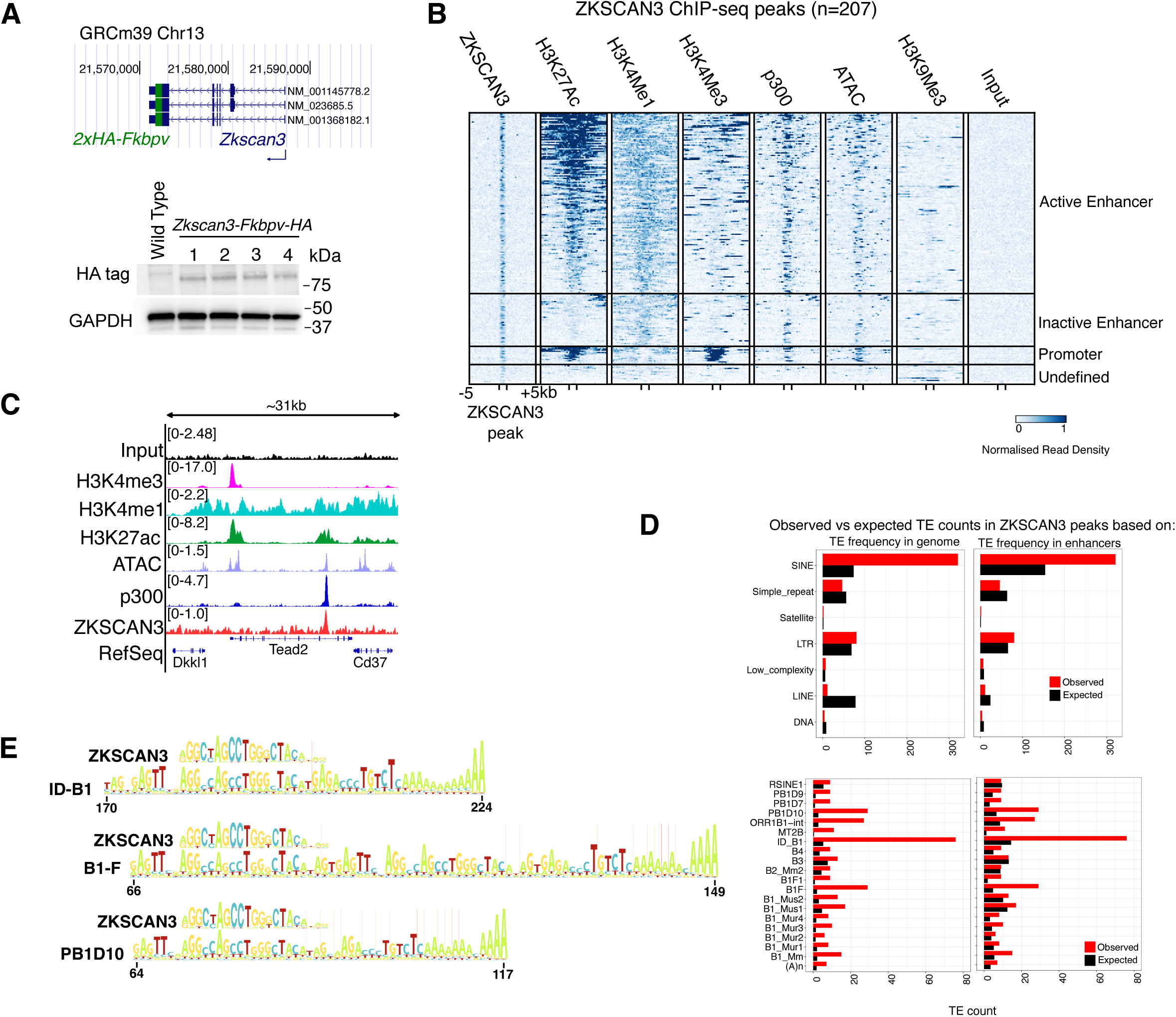
ZKSCAN3 binds to a SINE-embedded sequence within enhancers. **(A)** Schematic showing the tagging of endogenous *Zkscan3* with FKBPv and double HA tags, and immunoblot showing expression of ZKSCAN3-FKBPv-HA protein in four independent clonal lines. Annotated *Zkscan3* genes from NCBI RefSeq (curated subset) are shown. A non-specific band at ∼100kDa is also detected. **(B)** Heatmap showing normalised read density of ZKSCAN3, H3K27ac, H3K4me1, H3K4me3, p300 and H3K9me3 ChIP-seq and ATAC-seq signal over ZKSCAN3 peaks. ZKSCAN3 peaks are scaled to 1000bp and heatmaps show 5kb up and downstream for context. ZKSCAN3 peaks were partitioned into four groups based on intersection with ChIP-seq peaks of relevant histone modifications: active enhancers (H3K4me1 + H3K27ac), inactive enhancers (only H3K4me1), promoters (H3K4me3) and undefined regions (no intersection). ZKSCAN3 peaks are ordered by the level of H3K27ac. **(C)** Genome browser track example of a ZKSCAN3 ChIP-seq peak intersecting H3K4me1, H3K27ac and p300 ChIP-seq and ATAC-seq peaks at a putative enhancer. Tracks show the average of 4 (for ZKSCAN3) or 2 replicates. **(D)** Bar graphs showing the observed count of TEs, grouped by family (above) or sub-family (below), in ZKSCAN3 peaks compared with the expected count based on the background prevalence of TEs in the genome (left) or enhancers (right). Only sub-families with an observed count >5 are shown. **(E)** Alignment of the ZKSCAN3 binding motif to the consensus profile of the top three enriched SINE elements from (D). Only the 3’ end of the SINE elements are shown. Consensus models were sourced from Dfam(REF). Vertical lines in consensus models indicate sites of insertions, with darker colours indicating the frequency.

We used chromatin immunoprecipitation followed by sequencing (ChIP-seq) to assess ZKSCAN3 genomic binding locations and their chromatin context in ‘*Zkscan3-Fkbpv-HA’* mESCs. We detected 207 ZKSCAN3 peaks, the majority of which were marked with the chromatin signature of active enhancers (Figure 1B,C). 178/207 ZKSCAN3 peaks intersected a peak for enhancer-associated H3K4me1^34^, and 139 of these also intersected a peak for H3K27ac, which distinguishes active enhancers^35^. We also observed a small number of peaks (15/207) intersecting the promoter-associated histone mark H3K4me3, and a small number which had no characteristic chromatin signature (14/207).

KZFPs are generally associated with the recruitment of SETDB1 and the deposition of H3K9me3^36,37^. However, we observed no intersection of ZKSCAN3 peaks with H3K9me3-marked regions (Figure 1B). To corroborate this finding, we also assessed the impact of genetic *Zkscan3* ablation on H3K9me3, by generating *Zkscan3* knockout (*Zkscan3-KO*) mESCs (Figure S4). We profiled *Zkscan3-KO* mESCs cultured in both Serum/LIF (containing Foetal Bovine Serum, FBS, and Leukaemia Inhibitory Factor, LIF) and 2i/LIF (lacking FBS and containing LIF and inhibitors against the kinases Glycogen Synthase Kinase 3, GSK3, and Mitogen-activated protein kinase, MEK), conditions that recapitulate distinct stages of pluripotency^38–41^ (Figure S5). Using ChIP-seq we observed that the general distribution of H3K9me3 was unaffected, as was the total level of H3K9me3 as assessed by immunoblot (Figure S5A-C). Calling differentially enriched H3K9me3 peaks between Wild Type (WT) and *Zkscan3-KO* cells, we observed no significant changes in Serum/LIF-cultured cells and two significant changes in 2i/LIF-cultured cells, although neither of these intersected a ZKSCAN3 binding site (Figure S5D). Thus, ZKSCAN3 does not bind to heterochromatic regions of H3K9me3, nor does *Zkscan3* knockout result in changes to the deposition of H3K9me3 at its binding loci.

KZFPs are generally known to bind TEs^1,3,28,42,43^. To understand if ZKSCAN3 targets a particular class of TE, we calculated which TEs were overrepresented in ZKSCAN3 peaks. We observed that SINE retrotransposons were enriched over 4-fold at ZKSCAN3 binding sites compared with the genome as a whole, and over 2-fold compared with an unbiased sampling of other enhancer sequences (Figure 1D). Within the SINE family, certain sub-families were further enriched, including ID_B1 (76 observed vs 14 expected) and various B1 and Proto-B1 (PB1) SINEs (including B1_Mm, B1_Mus2, B1_Mus1, B1_Mur4, B1_Mur3, B1_Mur2, B1_Mur1, B1F, B1F1, PB1D7, PB1D10 and PB1D9; 162 observed vs 63 expected). B1, PB1 and ID_BI SINEs all contain highly related 3’ sequences and a common origin in 7SL-RNA: B1 and PB1 SINEs originate from the 7SL RNA^44–46^, while ID_B1 is a dimeric SINE formed from a fusion between a tRNA-derived sequence (at its 5’ end) and a 7SL RNA sequence (at its 3’ end)^46^.

We performed *de novo* motif analysis on the DNA sequence extracted from ZKSCAN3 peaks and identified a ZKSCAN3 binding motif present in all 207 peaks (E-value: 1.4e-846) (Figure 1E). Comparing this motif to the consensus profiles^47,48^ of the three most enriched SINE sub-families (ID_BI, B1F & PB1D10) revealed that the ZKSCAN3 binding motif matched a sequence located at the 3’ end of these elements (Figure 1E). Furthermore, a majority (150/207) of ZKSCAN3 peaks intersected a B1, PB1 or ID_B1 SINE, as annotated by RepeatMasker^49^. It is possible that a SINE sequence was not identified at the other ZKSCAN3 binding sites because of mutational drift. Indeed, others have suggested that over time, as TEs undergo mutational drift, all that may remain of a TE is the KZFP binding region, which undergoes positive selection due to its co-option as a regulatory region^3,4^. In sum, these data demonstrate that ZKSCAN3 binds to a motif embedded within the 3’ end of 7SL-RNA derived SINE retrotransposons.

To predict which TFs may co-localise with ZKSCAN3, we performed motif discovery for known TF binding motifs in ZKSCAN3 peak sequences using the HOCOMOCO Mouse (v11 CORE) database^50^. We observed significant enrichment for four nuclear receptors: Retinoid X Receptor Alpha (RXRA), Retinoic Acid Receptor Alpha (RARA), Estrogen Related Receptor B (ESRRB) and Nuclear Receptor Sub-family 5 Group A Member 2 (NR5A2) (Figure S6A). RARA and RXRA retinoic acid receptors heterodimerise to form a transcriptional regulator that is required for the development of many organs and tissues^51–56^. ESRRB and NR5A2 are orphan receptors that bind to highly similar motifs, co-occupy most of their binding loci, and are essential for the maintenance of the murine naïve pluripotent state, as well as the development of diverse tissues including the pancreas, jaw and brain^57–61^. Using publicly available ChIP-seq data for 2i/LIF-cultured mESCs, we determined that ESRRB occupied the majority of ZKSCAN3 peaks (142/207) (Figure S6B). Regions bound by ESRRB and NR5A2 are often co-bound by other pluripotency factors, including OCT4, SOX2 and NANOG^61,62^. Again, using publicly available ChIP-seq data for 2i/LIF-cultured mESCs, we found that ZKSCAN3 occupied many common sites with SOX2 (121/207), OCT4 (98/207) and NANOG (110/207) (Figure S6C). Thus, ZKSCAN3 co-occupies many sites with core pluripotency transcription factors, indicating potential co-regulation.

In summary, in mESCs ZKSCAN3 targets 7SL-RNA-derived SINE retrotransposons but its binding is not associated with H3K9me3 induction. Rather, its binding sites exhibit the epigenetic signature of enhancers and are co-occupied by pluripotency-associated transcription factors.

### ZKSCAN3 restricts accessibility and H3K27Ac at enhancers in mESCs

To explore the role of ZKSCAN3 in chromatin regulation we performed ChIP-seq for enhancer-associated histone marks H3K4me1 & H3K27Ac, promoter-associated mark H3K4me3 and chromatin accessibility (assessed by ATAC-seq) before and after degron-mediated ZKSCAN3 depletion. We treated two clonal *Zkscan3-Fkbpv-HA* cell lines with 100nM dTAG for 24hrs and confirmed the degradation of endogenous ZKSCAN3 by immunoblot (Figure 2A). Degradation was reversible and we were able to rescue ZKSCAN3 protein levels by removing dTAG from the medium for 5 days. Upon ZKSCAN3 degradation, we observed minimal changes to H3K4me1 and H3K4me3 (Figure S7A) but found that a subset of ZKSCAN3 binding sites gained H3K27ac and became more accessible (Figure 2B). We noted significant changes in chromatin accessibility at 32/207 binding sites (31 up, 1 down) and significant changes in the H3K7ac enrichment at 33/207 binding sites (33 up, 0 down) (limma, p<0.05). To understand the extent to which these gains in H3K27Ac enrichment and chromatin accessibility co-occurred, we binned ZKSCAN3 binding sites into those that gained both significant enrichment of H3K27Ac and accessibility (15/207), gained H3K27Ac only, (18/207) gained accessibility only (16/207) or showed no change in either (158/207) (Figure 2C). There were many instances of loci gaining H3K27Ac without a concurrent increase in accessibility and vice versa (Figure 2D). Therefore, the effects of ZKSCAN3 degradation were locus dependent. Using the categories we defined earlier (Figure 1B), we also observed that gains in H3K27Ac and accessibility occurred at ZKSCAN3 binding sites annotated as both active and inactive enhancers (Figure S7B,C). We concluded that ZKSCAN3 was not acting as a complete repressor, but exhibiting a rheostat-like behaviour, dampening rather than ablating enhancer accessibility and acetylation. Next, we rescued ZKSCAN3 by removing dTAG from the culture medium and observed that gains in H3K27Ac and accessibility were largely reversed (Figure 2E). At many loci ZKSCAN3 rescue restored the epigenetic state present in untreated cells (Figure 2F). These data therefore demonstrate a causal relationship between the binding of ZKSCAN3 and a reduction in the levels of H3K27Ac and chromatin accessibility at a subset of its binding sites. We therefore hypothesised that ZKSCAN3 limits the ability of enhancers to drive transcription of target genes.

**Figure 2.**
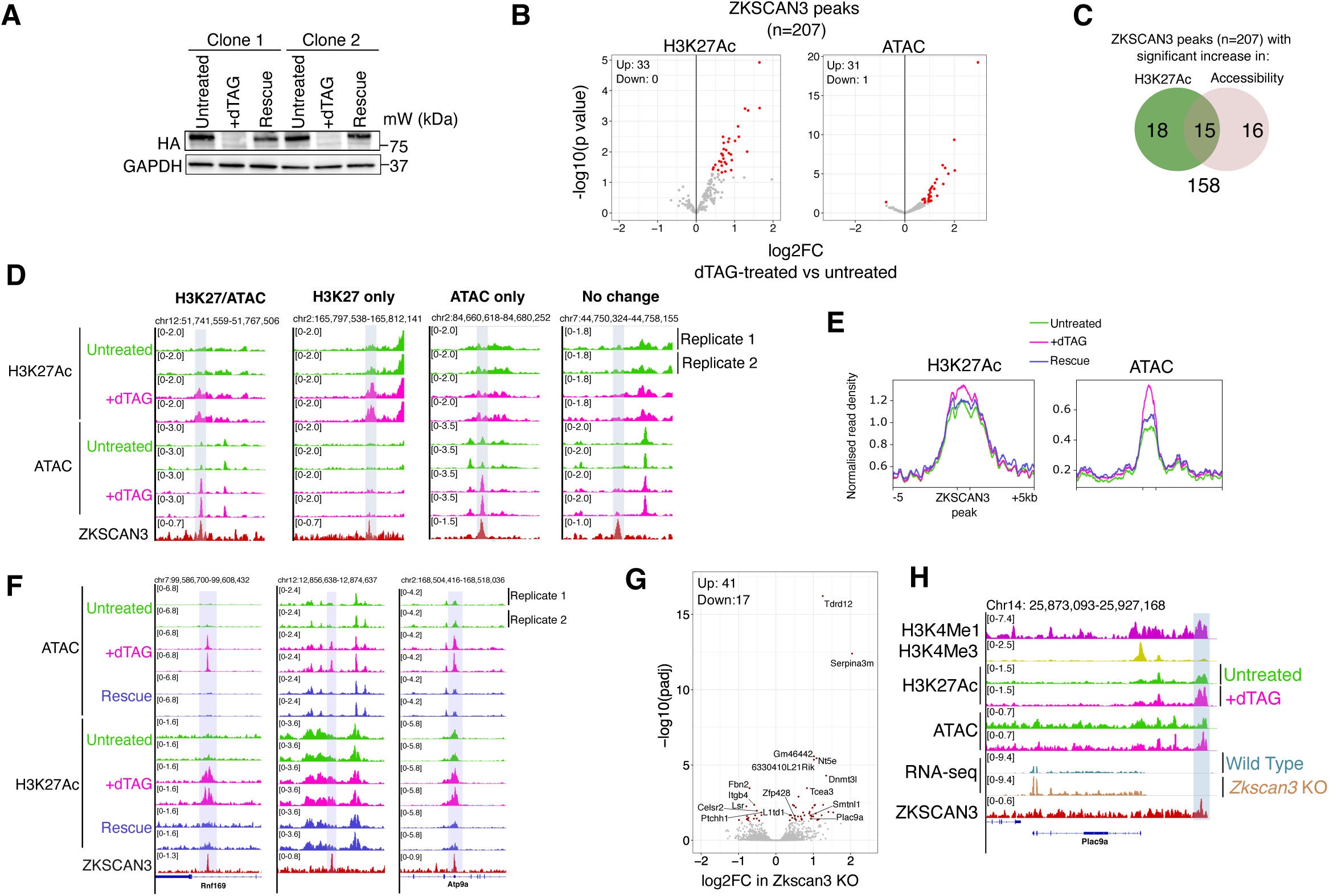
ZKSCAN3 restricts the accessibility and H3K27 acetylation of enhancers in mESCs. **(A)** Immunoblot showing degradation of ZKSCAN3-FKBPV-HA following treatment with 100nM dTAG for 24 hours and rescue of expression following removal of dTAG for 5 days. Experiments were performed in duplicate using independently derived clonal lines. **(B)** Volcano plots showing the change in H3K27ac enrichment (by ChIP-seq) and accessibility (by ATAC-seq) at ZKSCAN3 peaks in dTAG-treated cells. Significant hits are shown in red (p<0.05). **(C)** Venn diagram showing the number of ZKSCAN3 peaks that gained significant enrichment of H3K27Ac ChIP-seq or ATAC-seq signal, or gained no enrichment. **(D)** Genome browser track examples of different effects of ZKSCAN3 degradation at different loci. **(E)** Metaplot profiles showing the normalised read density of H3K27Ac ChIP-seq and ATAC-seq reads over ZKSCAN3 peaks in untreated, dTAG-treated and rescue cells. ZKSCAN3 peaks are scaled to 1000bp and the region 5kb up and downstream is shown for context. **(F)** Genome browser track examples of three loci where ZKSCAN3 degradation increased the accessibility and H3K27Ac levels at a ZKSCAN3 peak, and where rescue of ZKSCAN3 expression reversed the increase. **(G)** Volcano plot showing gene expression changes between WT and *Zkscan3*-KO mESCs. Significantly differentially expressed genes (p<0.05) are shown in red. **(H)** Genome browser track example of transcriptional changes to *Plac9a*, which is proximal to a ZKSCAN3-regulated putative enhancer. In all panels, profiles and genome browser tracks show the average signal of 4 replicates (ZKSCAN3 ChIP-seq), 3 replicates (RNA-seq) or 2 replicates (ChIP and ATAC-seq).

To investigate the impact of altered chromatin state in the absence of ZKSCAN3 on gene expression, we performed bulk RNA-seq in *Zkscan3-KO* mESCs and observed 58 differentially expressed genes (DEGs) (41 up, 17 down) (Figure 2G, Table S1). 17% of upregulated genes, but 0% of the downregulated genes, were immediately proximal (<50kb) to a ZKSCAN3 peak detected in WT cells, consistent with the hypothesis that ZKSCAN3 functions as a repressor. For example, *Plac9a*, which is proximal to a ZKSCAN3-occupied putative enhancer that gains both H3K27Ac and accessibility upon ZKSCAN3 degradation, is up-regulated in *Zkscan3-KO* mESCs (Figure 2H). Other examples included enhancers proximal to upregulated genes *Smtnl1* and *Csrp1* (Figure S8A-B). We quantified RNA-seq read density directly under and within 5kb of ZKSCAN3 peaks and observed that 19 peaks exhibited significantly higher read density in *Zkscan3-KO* cells, including reads mapping to unannotated, apparent non-coding transcripts (Figure S8C-E). Only one ZKSCAN3 peak exhibited a lower RNA-seq read density after *Zkscan3* ablation. In sum, these data suggest that ZKSCAN3 acts as a transcriptional repressor of both genic and non-genic transcripts by dampening the activity of enhancers in mESCs.

### ZKSCAN3-mediated repression is independent of its KRAB domain and KAP1

The lack of association between ZKSCAN3 function and heterochromatin prompted us to investigate the function of its KRAB domain. The KRAB domain consists of 50-75 amino acid residues divided into A and B boxes, of which the A box is essential to mediate transcriptional repression, and the B box enhances repression^42,63^. A multiple sequence alignment of ZKSCAN3 with 59 KRAB A boxes listed on InterPro^64^ reveals mutations at several highly conserved residues (Figure S9A). Importantly, these residues were previously identified as essential for both KAP1 recruitment and transcriptional repression (Figure 3A). Work from Peng and colleagues^65^ identified the residues essential for KAP1 recruitment in KRAB-O, a protein with a typical KRAB A-box. Using the BLOSUM62 matrix^66^ to define conservative amino acid substitutions, we found that the ZKSCAN3 KRAB A-box contains 3 non-conservative substitutions at residues identified as essential for KAP1 binding. Furthermore, Tycko and colleagues^67^ identified 12 residues in the ZNF10 KRAB A box that are essential for transcriptional repression and which they propose form a binding surface with KAP1. The ZKSCAN3 KRAB A box has non-conservative substitutions at 3 of these residues. Thus, ZKSCAN3 possesses a variant KRAB domain with several mutations that make it unlikely to recruit KAP1. Consistent with these predictions, we found that immunoprecipitation of exogenously expressed ZKSCAN3 did not identify KAP1 as an interactor, while KAP1 was readily immunoprecipitated with ZFP809, an established KAP1-interacting KZFP^14,68,69^, in the same assay (Figure 3B).

**Figure 3.**
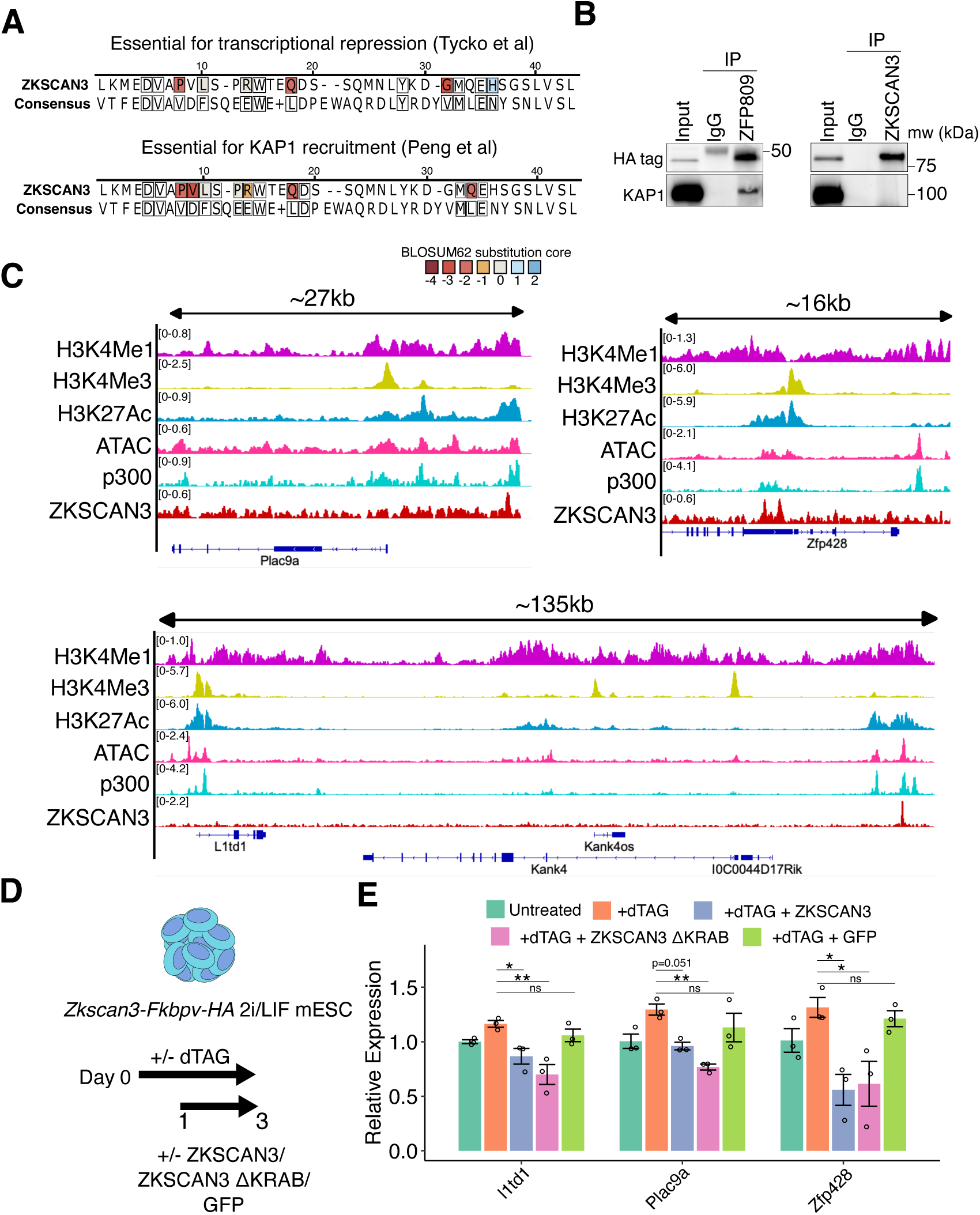
ZKSCAN3 repression is independent of the KRAB domain and KAP1. **(A)** Alignment of the ZKSCAN3 KRAB domain A-box to the A-box consensus sequence, derived from 59 KRAB A boxes listed on InterPro (Figure S9A). Residues identified as essential for KAP1 recruitment^67^ or transcriptional repression^69^ are identified with a black border. Substitutions in the ZKSCAN3 sequence are coloured by their BLOSUM62 substitution score, where blue shading represents a more conservative substitution and red more non-conservative. **(B)** Immunoblots showing co-immunoprecipitation of KAP1 with ZFP809, but not with ZKSCAN3. **(C)** Genome browser track images showing ZKSCAN3 binding sites proximal to *Plac9a*, *Zfp428* and *L1td1*. Tracks show the average of 4 (ZKSCAN3 ChIP-seq) or 2 replicates. **(D )** Schematic outlining experimental design for rescue of ZKSCAN3-mediated gene repression by transfection of different vectors. **(E)** Bar graphs showing the relative expression of *L1td1*, *Plac9a* and *Zfp428*. Bar graphs are plotted as the mean and error bars show SEM. Expression is presented as relative to untreated cells. Expression was normalised to the geometric mean of *Atp5b*, *Ubqc* and *Hprt*. Significance was assessed by one-way ANOVA followed by Tukey’s post-hoc test. *p<0.05, ** p <0.005, ns = not significant

To assess if the KRAB domain is necessary for ZKSCAN3-mediated transcriptional repression, we generated a mammalian expression vector containing a ZKSCAN3 mutant lacking the KRAB domain (ZKSCAN3 ΔKRAB), with a deletion of amino acid residues 213-273 (Figure S9B). We identified three genes that were up-regulated in *Zkscan3-KO* mESCs (*L1Td1*, *Plac9a* and *Zfp428*) and were proximal to a ZKSCAN3 peak, suggesting direct regulation (Figure 3C). We degraded ZKSCAN3 using dTAG and assessed whether introduction of WT ZKSCAN3, ZKSCAN3 ΔKRAB, or GFP could rescue gene repression (Figure 3D). In three independent *Zkscan3-Fkbpv-HA* clonal lines treated with dTAG to acutely deplete ZKSCAN3, *L1Td1*, *Plac9a* and *Zfp428* were up-regulated, reproducing results observed in *Zkscan3-KO* cells (Figure 3E). The expression of these genes was rescued in dTAG-treated cells by exogenous ZKSCAN3, with all three genes showing a reduction to untreated cell levels, or even below (for *Zfp428*), likely owing to higher expression of the exogenous gene. Exogenous expression of GFP failed to rescue the dTAG-mediated changes. Notably, ZKSCAN3 ΔKRAB was able to rescue expression of all three genes, similarly to full length ZKSCAN3 (Figure 3E). These results indicate that the KRAB domain is not required for ZKSCAN3-mediated repression. Together these data suggest that ZKSCAN3 represses enhancer activity through a mechanism independent of KRAB-mediated KAP1 recruitment.

### Chromatin-bound ZKSCAN3 associates with transcriptional regulators of neural development

To further interrogate ZKSCAN3-occupied sites, we performed Rapid Immunoprecipitation of Endogenous Proteins (RIME), a cross-linking mass spectrometry-based method optimised for the identification of proteins within the immediate environment around chromatin-bound transcription factors^70^. dTAG-treated cells, where endogenous ZKSCAN3 was degraded, were used as a negative control for these experiments, allowing us to compare pull-downs with the same anti-HA antibody.

We identified 34 proteins that were significantly enriched in untreated relative to ZKSCAN3-degraded cells (Figure 4A and Table S2). As expected, the two most enriched proteins were ZKSCAN3 itself and FKBP1A (arising from peptides mapping to the FKBPv degron tag). Using the STRING database^71^, we observed that many of the other significantly enriched proteins were previously reported to interact or exist in complexes (Figure 4B). The significant hits consisted of mainly transcription factors and associated co-regulators. The most significant GO term for the proteins enriched in RIME was “transcription regulator complex”, with other significant terms including “protein-DNA complex” and “transcription factor binding” (Figure 4C). Consistent with ZKSCAN3 localisation at enhancers, we observed significant enrichment of enhancer-associated acetyltransferase p300 and the H3K4 demethylase KDM1A, which occupies most enhancers in mESCs^72^. The transcription factor ESRRB, which our earlier analysis found to be bound at many ZKSCAN3 binding sites (Figure S6), was also significantly enriched. Notably, KAP1 was not enriched, providing further evidence that ZKSCAN3 acts in a KAP1-independent manner.

**Figure 4.**
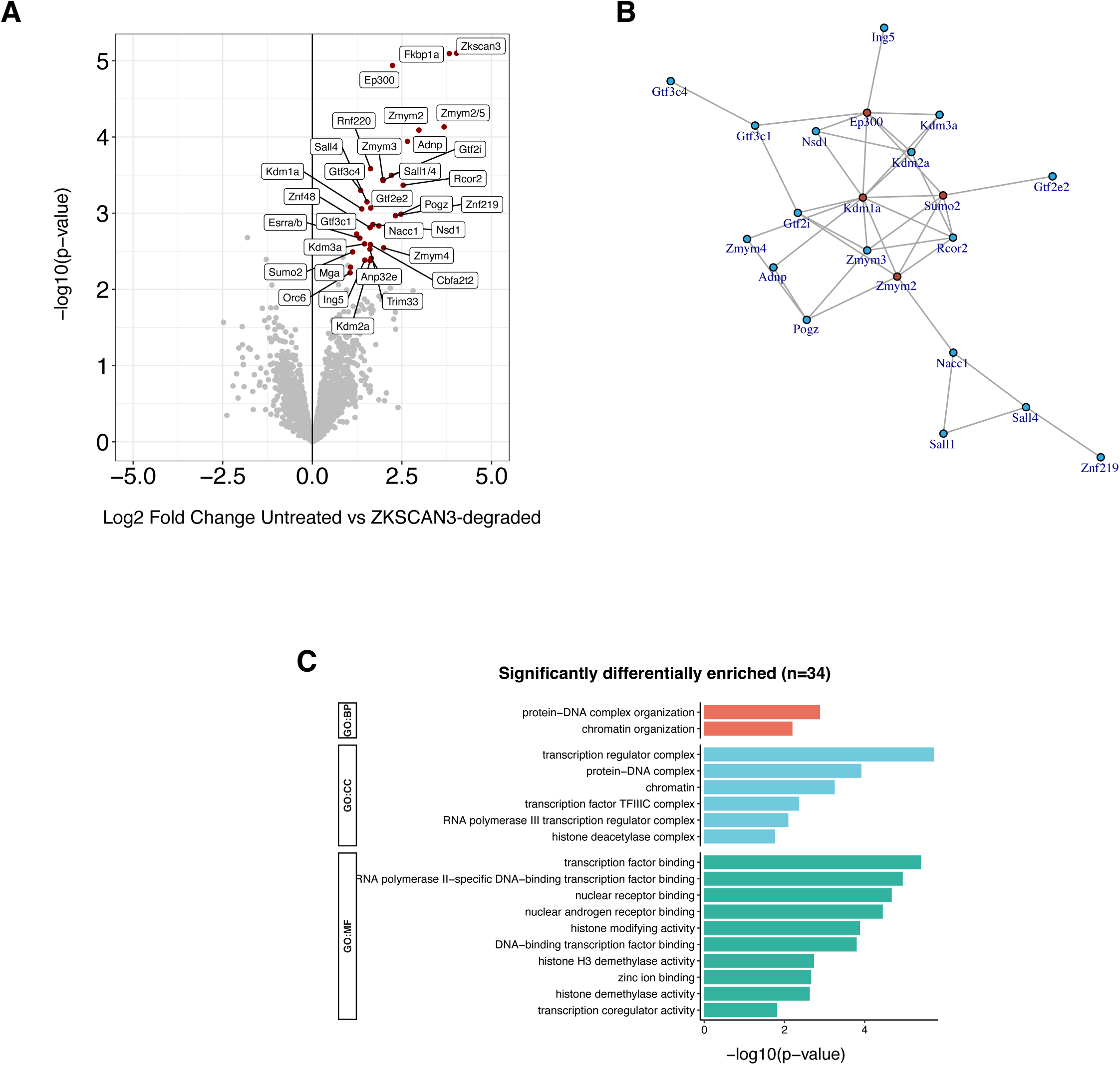
Chromatin-bound ZKSCAN3 localises with transcriptional regulators of neural development. **(A)** Volcano plot showing the enrichment of proteins detected in ZKSCAN3 RIME pulldowns between control and ZKSCAN3-degraded cells. Significantly differentially enriched proteins (LFC>1, p<0.05), are shown in red and labelled. **(B)** Network of putative protein interactions between significantly differentially enriched proteins from the STRING database. Only interactions with a score >300 are shown. Proteins with >6 interactions are shown in red. **(C)** Gene Ontology (GO) analysis of significantly enriched proteins. Only GO terms with <1000 genes are shown. CC = Cellular Component; MF = Molecular Function; BP = Biological Process.

Many of the enriched proteins are transcriptional regulators involved in neural differentiation, or are implicated in neural differentiation by being genetically associated with human neurodevelopmental disorders (NDDs) (Table 1). To take several key examples, the transcriptional regulators *Adnp* and *Pogz* are the two most commonly mutated genes in autism spectrum disorder^73–77^; mutations to the transcription factor *Gtf2i* are the main genetic cause of altered neurocognitive profile in Williams-Beuren syndrome^78–80^; and the TF *Zmym3* is associated with SZ and bipolar disorder risk^81,82^. Furthermore, cross-referencing our RIME hits with SysNDD, a curated database of gene-disease relationships in NDDs^83^, revealed that 41% of the RIME-identified regulators were previously implicated in NDD, implying a role in neural development (Table S3). Perturbation of *Adnp*, *Pogz* or *Gtf2i* in pluripotent cells causes defects in neural differentiation^84–88^, suggesting their activity in pluripotent cells facilitates later neural differentiation events. Given the association of ZKSCAN3 with neural TFs, and the previously reported genetic associations with SZ in humans, we hypothesised that ZKSCAN3 itself may play a role in neural differentiation.

**Table 1.** ZKSCAN3 localises with transcriptional regulators associated with neural differentiation and neurodevelopmental disorders.

| Protein | Function | Laboratory work indicating neural role | Human genetic pathology |
| --- | --- | --- | --- |
| ADNP | Transcription factor associated with chromatin re-modelling<sup>163,164</sup> | *Adnp* knockout in mESCs results in failure to differentiate to the neuronal lineage<sup>84</sup>. *Adnp* knockout mice are not viable, due to lethal defects in brain formation and neural tube closure<sup>165</sup>. | Autism, Intellectual Disability<sup>73–76</sup> |
| POGZ | Transcriptional co-regulator<sup>85,166</sup> | *Pogz* knockout causes embryonic lethality<sup>167</sup>, whereas *Pogz* heterozygous knockouts show growth delay, smaller brain size, defects in neurogenesis and autism-like behaviour<sup>168,169</sup>. *Pogz* knockdown inhibits proper neuronal development of mESCs and causes defects in cultured mouse cortical neurons<sup>85,86</sup>. Mechanistically, POGZ controls chromatin accessibility at enhancers and promoters and promotes neuronal gene expression<sup>170</sup>. | Autism, intellectual disability<sup>171–177</sup> |
| GTF2I | Transcriptional repressor | GTF2I knockout mice are not viable, while heterozygous mice exhibit microcephaly, cranio-facial alterations, altered axonal outgrowth and altered social behaviour<sup>178–181</sup>. Conditional knockout in neurons results in reduced numbers of oligodendrocytes, reduced myelin thickness and diminished conductivity of axons<sup>87</sup>. GTF2i KO hESCs differentiated to neurons have altered electrochemical physiology and exhibit synaptic dysfunction<sup>88</sup>. | 7q11.23 dosage dependent neurodevelopmental disorders; deletion causes Williams-Beuren syndrome (hyper-sociability, intellectual disability, cranio-facial abnormalities) while duplication frequently results in autism spectrum disorder<sup>78–80</sup>. |
| RNF220 | E3 ubiquitin ligase.<sup>182</sup> | Required for proper development of oligodendrocytes and noradrenergic neurons and is essential for cerebellum development<sup>183–186</sup>. *Rnf220* knockout in NPCs inhibits maintenance and promotes differentiation<sup>187</sup>. Regulates the patterning of the ventral spinal neural tube in mice<sup>188</sup>. Regulates synaptic transmission through ubiquitination of synaptic receptors<sup>189</sup>. | Leukodystrophy; neurodegenerative disease with systemic implications<sup>190</sup> |
| NACC1 | Transcriptional co-repressor<sup>191–194</sup> | Facilitates proper differentiation by controlling a sub-network of pluripotency factors to repress neuroectodermal fate<sup>194</sup>. Required for proper glutamatergic transmission in mature neurons<sup>195</sup>. | Neurodevelopmental disorder including intellectual disability, microcephaly, and epilepsy<sup>196,197</sup> |
| ZMYM2 | Transcriptional repressor<sup>198</sup>. |  | Neurodevelopmental-craniofacial syndrome with variable renal and cardiac abnormalities<sup>199</sup> |
| ZMYM3 | Poorly characterised transcription factor. |  | X-linked intellectual disability (56,57). Variation in the number of Short Tandem Repeats (STRs) in the 5'UTR of ZMYM3 have been linked to schizophrenia and bipolar disorder risk<sup>81,82</sup>. |
| ZMYM4 | Poorly characterised transcription factor<sup>202,203</sup>. | Regulates expression of embryonic cranial genes critical for craniofacial development<sup>204</sup> |  |
| ZMYM5 | Poorly characterised transcription factor. Paralog of ZMYM2. |  |  |
| KDM1A | Lysine demethylase for H3K4 or H3K9<sup>205</sup>. Facilitates enhancer inactivation upon differentiation<sup>206</sup>. | KDM1A knockout mice are not viable<sup>207</sup>; cKO in Nestin positive cells exhibit motor defects and abnormal brain morphology<sup>208</sup>. KDM1A is required for maintenance and proliferation of neural progenitors and their proper differentiation to mature cortical neurons<sup>209–213</sup>. | Developmental disorder which includes cognitive impairment and developmental delay<sup>214,215</sup>. |
| KDM2A | Lysine demethylase for H3K36<sup>216</sup> | In hNPCs, *KDM2A* knockout impairs proliferation, increases apoptosis, induces premature neuronal differentiation, and, upon differentiation to mature neurons, affects synapse maturation<sup>217</sup>. | Intellectual disability<sup>217</sup> |
| KDM3A | Lysine demethylase for H3K9Me1/2 <sup>218</sup> | Essential for primary neuron formation in *Xenopus*; knockdown causes abnormal neural crest migration and inappropriate expression of neural markers <sup>219,220</sup> |  |
| SALL4 | Transcription factor, which can act as a repressor or activator dependent upon context <sup>221,222</sup>. Key regulator of the pluripotent state <sup>223–225</sup>. | Facilitates neural differentiation by preventing precocious activation of neural genes in mESCs <sup>226</sup>. |  |
| RCOR2 | Transcriptional co-repressor <sup>227</sup> | Orchestrates neurogenesis in the developing mouse brain; conditional knockout mice displayed reduced brain sizes and cortical thickness, as well as structural irregularities in the brain layers, a decrease in the number of neuronal progenitors and neurons, and an increase in cell death. <sup>227,228</sup> |  |
| GTFIIIC1/4 | Components of the GTFIIIC transcription factor complex. Required for Pol III mediated transcription, but also has Pol III independent functions <sup>229,230</sup>. Possesses HAT activity <sup>231</sup> | In hESCs, GTFIIIC is bound at loci associated with neuronal differentiation and co-occupies many ADNP sites <sup>232</sup>. In mature cultured cortical neurons, perturbation affects dendritic length and branching <sup>233</sup> |  |
| ZNF219 | Transcriptional repressor <sup>234–236</sup> |  |  |
| ZNF48 | Poorly characterised transcription factor. |  |  |
| Cbfa2t2 | Transcriptional co-repressor<sup>237</sup> |  |  |
| ANP32E | Histone chaperone facilitating exchange of H2AZ from the nucleosome, especially from enhancers<sup>238,239</sup> |  |  |
| ING5 | Putative transcriptional co-regulator which is a reader protein of H3K4Me3 and essential for H3K14Ac deposition<sup>240</sup> |  |  |
| MGA | Transcriptional repressor associated with non-canonical Polycomb complex 1.6<sup>241,242</sup> |  |  |
| Rbm33 | Poorly characterised RNA binding protein<sup>243</sup> |  |  |
| GTF2E2 | Component of the TFIIE complex required for RNA Pol II mediated transcription<sup>244</sup> |  |  |
| NSD1 | H3K36 methyltransferase<sup>245,246</sup> | Required for establishment and maintenance of neuronal identities in mice<sup>246</sup> | Sotos syndrome; developmental disorder characterised by gigantism, craniofacial abnormalities, and frequently autism spectrum disorder<sup>247–249</sup> |
| EP300 | Enhancer-associated histone acetylase<sup>250–252</sup> |  |  |
| ESRRB | Transcription factor associated with maintenance of naïve pluripotency<sup>61,62,253,254</sup> |  |  |
| Trim33 | Transcriptional regulator with chromatin reader and E3 ubiquitin ligase functions<sup>255–258</sup> |  |  |
| SUMO2 | Small ubiquitin-like modifier which can be attached to proteins to modulate function<sup>259,260</sup> |  |  |
| Pkp4 | Poorly characterised protein involved in cell-adhesion<sup>261</sup> |  |  |
| Orc6 | Component of the origin recognition complex (ORC) that binds origins of replication<sup>262,263</sup> |  |  |

### ZKSCAN3 regulates enhancers during neural differentiation

To investigate the role of ZKSCAN3 in neural differentiation, we cultured *Zkscan3-Fkbpv-HA* mESCs in N2B27 medium without LIF for 6 days, following an established method for neural differentiation^89–93^. We performed ChIP-seq for ZKSCAN3 in untreated neural cells, and RNA-seq and ChIP-seq for H3K27Ac, H3K4me1 and H3K4me3 in both untreated and ZKSCAN3-degraded cells (Figure 5A). dTAG was added to mESCs 24 hours prior to the start of the differentiation protocol, to degrade ZKSCAN3 before the onset of differentiation, and then supplemented into the medium daily. We confirmed that the dTAG molecule alone did not affect the ability of WT cells to differentiate, as assessed by the induction of neural progenitor markers *Nestin* and *Pax6* and downregulation of the pluripotency marker *Pou5f1* (Figure S10).

**Figure 5.**
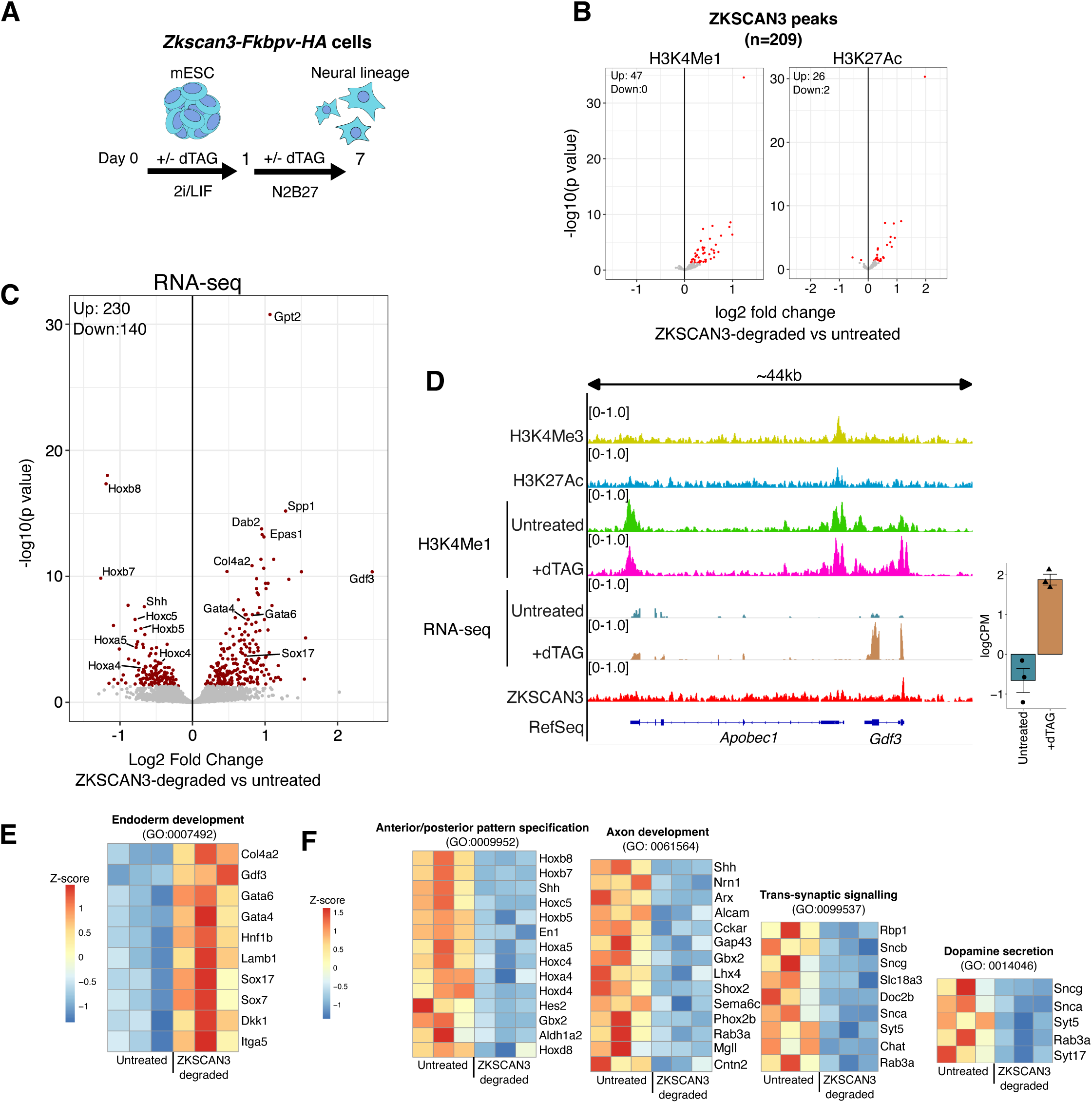
ZKSCAN3 is a repressor of enhancers during neural differentiation. **(A)** Schematic outlining experimental design for neural differentiation of mESCs. **(B)** Volcano plot showing differential enrichment of H3K4me1 and H3K27ac at ZKSCAN3 peaks in untreated and dTAG-treated cells upon neural differentiation. Significant hits are shown in red (p<0.05). **(C)** Volcano plot showing significant gene expression changes, as measured by bulk RNA-seq, in ZKSCAN3-degraded neural lineage cells. Significant differentially expressed genes (p<0.05) are shown in red. **(D)** Genome browser track example showing the direct targeting of *Gdf3* by ZKSCAN3. Tracks show the average signal of 3 replicates (RNA-seq) or 2 replicates (ChIP and ATAC-seq). For clarity, only RNA-seq reads in the direction of the upregulated gene are shown. A bar chart showing the full RNA-seq data for all three replicates is shown on the right. **(E)** Heatmap showing gene expression changes for genes in the ‘endoderm development’ ontology. **(F)** As in (E), for various ontologies of up-regulated genes.

We identified 209 ZKSCAN3 binding sites in neural cells, all of which contained the same SINE-embedded binding motif we detected in mESCs (E value: 4.0e-517) (Figure S11A). Of these, 79 peaks also retained an identifiable 7SL RNA-derived SINE sequence in RepeatMasker annotation^49^. ZKSCAN3 was profoundly redistributed in neural cells compared to mESCs, with only 13 of the binding sites being the same between the two cell types (Figure S11B). Using ChIP-seq of enhancer-and promoter-associated marks to classify ZKSCAN3 peaks, we observed that in neural cells ZKSCAN3 intersected 83 H3K4me1^+^ /H3K27Ac^+^ peaks (putative active enhancers), 47 H3K4me1^+^/H3K27Ac^-^ peaks (putative primed enhancers), 69 H3K4me3^+^ peaks (putative promoters) and 10 regions devoid of either of these three marks (H3K4me1/3^-^/H3K27Ac^-^) (Figure S11C). In ZKSCAN3-degraded neural cells, a subset of ZKSCAN3 binding sites gained enrichment of H3K4me1 and H3K27Ac (Figure 5B). 47/209 ZKSCAN3 binding sites were differentially enriched for H3K4me1 (47 up, 0 down) and 28/209 sites were enriched for H3K27Ac (26 up, 2 down) (limma, p<0.05). We did not observe any significant changes to H3K4me3 at ZKSCAN3 peaks (Figure S11D). Therefore, in the neural lineage ZKSCAN3 localises to the same SINE-embedded motif and restricts the enrichment of enhancer-associated H3K4me1 and activation associated H3K27ac at its binding sites.

### ZKSCAN3 represses the endodermal and mesodermal fates and promotes the expression of axon guidance receptors during neural differentiation

To study the transcriptional effects of ZKSCAN3 degradation in neural cells, we performed bulk RNA-seq analysis. We identified 370 differentially expressed genes (230 up, 140 down) between ZKSCAN3-degraded and untreated cells (DESeq2, p<0.05) (Figure 5C, Table S5). The most upregulated gene was *Gdf3* (5.7 fold), a TF associated with the formation of mesoderm and endoderm^94,95^. The *Gdf3* locus is directly bound by ZKSCAN3, and this binding site gained enrichment of H3K4me1 upon ZKSCAN3 degradation (Figure 5D). Strikingly, in ZKSCAN3-degraded cells we also observed upregulation of key endodermal lineage markers *Gata4*, *Gata6* and *Sox17.* In total, nine genes in the “endoderm differentiation” gene ontology were significantly upregulated, including *Gdf3* and *Hnf1b* (Figure 5E), which were found to be direct targets of ZKSCAN3 in our ChIP-seq experiments.

Downregulated genes included seven Hox genes (*Hoxa4*, *Hoxc4*, *Hoxa5*, *Hoxb5*, *Hoxc5*, *Hoxb7* and *Hoxb8*) as well as other factors implicated in anterior-posterior specification including *Sonic hedgehog (Shh)* and the homeobox genes *En1* and *Gbx2* (Figure 5F). Other downregulated genes related to neural function, with significant enrichment of GO terms for “axonogenesis”, “trans-synaptic signalling” and “dopamine secretion” (Figure S12A). Accordingly, downregulated genes were significantly enriched in cellular components “axon”, “neuronal cell body” and “pre-synapse”. Finally, we cross-referenced our DEGs with the CellMarker 2.0 database^96^, which contains manually curated cell markers based on single cell RNA-sequencing (scRNA-seq) data and observed that upregulated genes most significantly included markers for primitive endoderm, whereas downregulated genes were most highly associated with brain motor neurons (Figure S12B). Together, these data indicate that ZKSCAN3 is responsible for regulating enhancers in the neural lineage, and that perturbation of ZKSCAN3 causes an aberrant gene expression signature characterised by up-regulation of endodermal markers and downregulation of genes associated with neural processes.

To gain a more complete view of the role of ZKSCAN3 during multi-lineage differentiation, we performed scRNA-seq of Embryoid Bodies (EBs) differentiated from *Zkscan3-Fkbpv-HA* cells with and without the presence of dTAG. We confirmed that the dTAG molecule alone did not affect the ability of WT cells to form EBs, induce ectodermal, mesodermal and endodermal lineage markers, or downregulate pluripotency markers within them (Figure S13). In scRNA-seq of *Zkscan3-Fkbpv-HA* cells differentiated to EBs, all the major cell clusters were present in both untreated and ZKSCAN3-degraded EBs (Figure S14A). *Zkscan3-*null mice are viable and born in Mendelian ratios^97^, and thus our findings are consistent with the idea that ZKSCAN3 is not necessary for the induction of a particular developmental lineage. In agreement with data from the Mouse Gastrulation Atlas, a publicly available scRNA-seq dataset of the early mouse embryo^98^, ZKSCAN3 was ubiquitously expressed in all lineages (Figure S14B-C). We used the Atlas to annotate the identity of the Seurat clusters in our EBs and were able to identify several cell types including endothelial cells, haematopoietic progenitors, erythrocytes, cardiomyocytes and neuroectodermal cells (Figure 6A).

**Figure 6.**
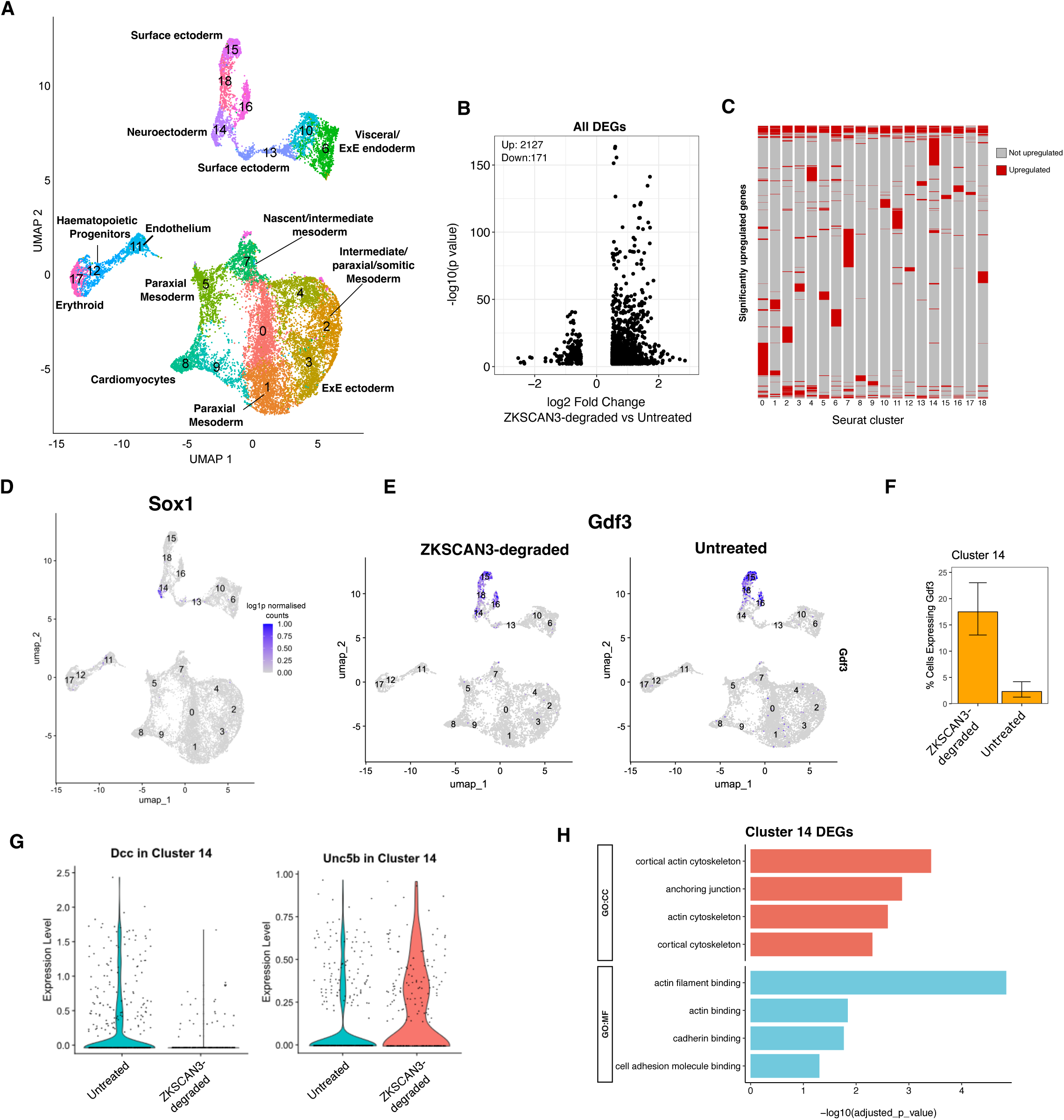
ZKSCAN3 regulates the expression of axon-guidance receptors. **(A)** UMAP projection of untreated Emybroid Bodies (EBs) and annotation of clusters based on the Mouse Gastrulation Atlas. Cells are coloured by Seurat cluster. **(B)** Volcano plot of scRNA-seq data showing the change in gene expression between untreated and ZKSCAN3-degraded EBs. **(C)** Heatmap showing the distribution of significant DEGs across clusters. Most clusters show distinct sets of upregulated genes. **(D)** UMAP projection of *Sox1* expression in untreated EBs, marking cluster 14 as neuroectoderm. **(E)** UMAP projection of *Gdf3* expression in untreated and ZKSCAN3-degraded EBs. **(F)** Bar chart showing the percentage of cells expressing *Gdf3* in cluster 14 in untreated and ZKSCAN3-degraded EBs. Error bars show 95% confidence intervals. **(G)** Violin plots showing expression of *Dcc* and *Unc5b* in untreated and ZKSCAN3-degraded neuroectoderm from EBs. **(H)** Gene ontology analysis of differentially expressed genes in cluster 14 (neuroectoderm) of ZKSCAN3-degraded EBs.

We identified 2127 upregulated and 171 downregulated genes across the clusters (p<0.05, L2FC>0.5) (Figure 6B). We first focussed on upregulated genes as, based on our previous characterisation of ZKSCAN3 acting as a repressor, they were more likely to represent direct targets. While upregulated genes were detected in all 18 clusters of the dataset (Figure 6C) consistent with the expression of *Zkscan3* across all clusters (Figure S14B-C), each cluster upregulated a largely distinct set of genes, with 780/1080 upregulated genes detected in a single cluster of cells, and a further 165/1080 detected in just two. These data suggest that ZKSCAN3 represses distinct sets of genes in different lineages.

GO analysis of all downregulated genes across clusters did not return any significant terms. In contrast, the upregulated genes were enriched for transcription factors (Figure S15A). We further analysed all upregulated TFs using WikiPathways and found significant enrichment for the mesodermal commitment pathway, pathways relating to the development of mesoderm-derived renal, cardiac and adipose cells, as well as endodermal commitment (Figure S15B). TFs in this category included master mesendodermal factors *Eomes* and *Brachyury,* which were upregulated within cells annotated as nascent/intermediate mesoderm and neuroectoderm (Figure S15C). Notably, ZKSCAN3 has been previously reported to regulate erythropoiesis^99^, and *Eomes*, which is a major regulator of this process^100^, was the most upregulated gene in both haematopoietic progenitors (8.6 fold) and erythrocytes (12.1 fold) in ZKSCAN3-depleted EBs. In sum, the genes repressed by ZKSCAN3 across lineages are enriched for transcription factors associated with the formation of mesoderm and endoderm.

To further assess the role of ZKSCAN3 in neural differentiation we focused specifically on cluster 14, which was annotated as neuroectoderm. Cluster 14 contained the only population of cells expressing *Sox1*, the earliest specific marker of neural precursors in the mouse embryo^101^ (Figure 6D). Within this cluster we observed upregulation of mesendodermal TF *Gdf3* (Figure 6E,F), mirroring our results in the bulk RNA-seq of neural lineage cells.

DCC (Netrin Receptor) and UNC5B (Netrin Receptor B) are cell-surface receptors that direct axonal navigation during neurodevelopment; Netrin binding to DCC results in attractive responses and Netrin binding to UNC5 (in some cases dimerised with DCC) results in repulsive responses^102–107^. *Dcc* was the most downregulated gene in Cluster 14 (L2FC = -2.4) (Figure 6G). Furthermore, we also observed significant upregulation of *Unc5b* (L2FC = 0.8) (Figure 6G). Netrin signalling facilitates remodelling of the actin cytoskeleton to direct the developing axon^102^. GO analysis of the genes differentially expressed in Cluster 14 returned a striking enrichment for actin-related terms: the most significant Molecular Function ascribed to these genes was “actin filament binding” and the most significant Cellular Component was “cortical actin cytoskeleton” (Figure 6H). Importantly, these actin-binding genes included *Cfl1* and *Dstn* (ADF), two proteins belonging to the ADF/cofilin family that are responsive to Netrin and act as key regulators of actin dynamics during neural differentiation^108–112^. We also observed upregulation of *Wdr1* (AIP1), the major cofactor of ADP/Cofilin^113^, as well as several other actin-binding proteins implicated in control of axonal growth including *Vcl* ^114^, *Tpm1* ^115^, *Cotl1* ^116^ and *Mical1* ^117–119^. In summary, ZKSCAN3 degradation results in changes to the expression of Netrin-responsive axon guidance receptors and their downstream effectors. Overall, the above findings indicate that ZKSCAN3 mediates proper gene expression during neural lineage commitment.

## Discussion

KRAB Zinc Finger proteins are well-established binders and repressors of transposable elements. In this study we report that the KZFP ZKSCAN3 controls gene expression by repressing enhancer activity, through binding to an enhancer-embedded SINE retrotransposon sequence. As such, our findings contribute to the emerging idea that a significant proportion of *cis*-regulatory sequences are derived from TEs^16,120–122^.

*Zkscan3* has homologs in most eutherians (placental mammals) but not in marsupials^123^, implying an emergence in the common ancestor of modern eutheria. 7SL RNA-derived SINEs are proposed to have emerged in either the common ancestor of Euarchontoglires^124^, or at an earlier time in the common ancestor of the Boreoeutherians^125^, although their origin is poorly understood. Both categories are in the eutherian clade, meaning that based on current evidence ZKSCAN3 emerged before 7SL-RNA-derived SINEs. One could speculate that, in such a scenario, SINE retrotransposons may have spread the existing ZKSCAN3 binding motif to new genomic locations, with the potential to re-wire gene regulatory networks by introducing ZKSCAN3 binding to new loci. This would follow numerous examples of transposon-mediated expansion of TF binding sites^126,127^, for example binding sites for OCT4 and NANOG in human embryonic stem cells^128^, interferon-responsive IRF1 and STAT1 in the context of innate immunity^129^, and a large number of TFs in the context of mammalian pregnancy^130^. Regardless of the exact mechanism of domestication, our findings contribute to an emerging understanding of how host-TE interactions shape gene expression networks.

To our knowledge, two examples of KZFP-mediated enhancer regulation have been described previously: firstly, over-expression of the KZFP ZNF611 in human naive ESCs induces induction of H3K9me3, loss of H3K27ac, and repression of hundreds of genes proximal to an evolutionarily young class of retrotransposons known as SVA elements^17^. Similarly, the KZFP ZNF676 recruits KAP1 to repress the apparent enhancer activity of LTR12C elements^131^. In these reports, KZFP-mediated enhancer regulation has been understood to occur by selective induction of heterochromatin. However, ZKSCAN3 belongs to the ZKSCAN sub-family of KZFPs that are less associated with KAP1 recruitment^3,28^. It has been suggested that the ancestral progenitor of ZKSCAN proteins functioned to repress TEs via KAP1, but that over time this sub-family has evolved KAP1-independent functions^7^. Here, we conclude that ZKSCAN3 does not recruit KAP1 owing to several pieces of converging evidence: (1) ZKSCAN3 has a mutated KRAB domain, including key residues associated with KAP1 recruitment; (2) ZKSCAN3 does not immunoprecipitate with KAP1; (3) KAP1 is not an interactor of ZKSCAN3 in a RIME mass-spectrometry screen; and (4) ZKSCAN3 can repress transcription when its KRAB domain is deleted. We find that rather than repressing enhancer activity via KAP1-mediated heterochromatin induction, the ZKSCAN protein ZKSCAN3 restricts H3K4me1, H3K27ac and chromatin accessibility at enhancer sites, and that this chromatin regulation is associated with reduced transcription of enhancer-proximal genes. This work provides, to our knowledge, the first functional study of a KZFP exhibiting such KAP1-independent activities.

We found that the effects of ZKSCAN3-mediated chromatin regulation are context-dependent, both in terms of the genomic locus and the cell type. Upon ZKSCAN3 degradation, an increase in the enrichment of H3K27ac at a ZKSCAN3 binding site was not necessarily associated with an increase in chromatin accessibility and vice versa. The enrichment of H3K4me1 was unaffected upon ZKSCAN3 degradation in pluripotent mESCs, but increased at some binding sites in cells undergoing neural differentiation. It is probable that, at a given genomic locus, the precise effects of ZKSCAN3 regulation are dependent on the specific chromatin microenvironment, consisting of the locus-specific and cell-type specific repertoire of TFs, co-regulators, and histone modifications. Our data emphasise the importance of considering chromatin regulation by TFs as a complex interplay between the mechanism of action of the TF and the existing chromatin environment at a given locus.

We observed ZKSCAN3 binding at both active and inactive enhancers (Figure 1B). Additionally, upon ZKSCAN3 degradation, increases in accessibility and H3K27ac occurred at both active and inactive enhancers (Figure S7). Based on our findings, we propose that ZKSCAN3 does not act as a complete repressor of enhancer activity, but exhibits a rheostat-like behaviour, dampening enhancer output to fine-tune the optimal expression level. Indeed, perturbation of ZKSCAN3-mediated enhancer regulation by protein degradation or knockout consistently resulted in mild transcriptional changes, with putative direct targets upregulated approximately 1.5-to 3-fold. This observation mirrors the findings of several previous publications using *Zkscan3* knockout or knockdown cells, where target genes were induced to a similar magnitude^99,132–134^. Therefore, data from this study and from others suggests that ZKSCAN3 does not inactivate transcription, but rather fine-tunes expression levels. In summary, we propose that ZKSCAN3 acts as a context-dependent rheostat, rather than an absolute repressor.

*Zkscan3-null* mice are viable, grossly phenotypically normal and born in Mendelian ratios^97^. Therefore, loss of ZKSCAN3 during cellular differentiation is unlikely to cause a dramatic phenotype such as failure to form a particular lineage. Rather, loss of ZKSCAN3-mediated enhancer regulation may result in inappropriate gene expression levels that manifest as subtler phenotypes, where the organism is grossly developmentally normal but exhibits pathology in certain systems. For example, *Zkscan3-null* mice exhibit a decreased number of late erythroblasts in the bone marrow^99^. ZKSCAN3 has been genetically associated with schizophrenia risk in humans. *ZKSCAN3* is located in a genomic region defined by the Psychiatric Genomics Consortium as the most associated with SZ risk^29^, and a SNP (rs733743) causing a missense variant in the *ZKSCAN3* coding sequence is significantly associated with SZ^30^. Additionally, 9 of the 39 SNPs that are significantly associated with both SZ and regional grey matter volume changes are *ZKSCAN3* eQTLs in adult brain tissue^31^. Our own analysis shows that many of these SNPs, or ones in tight linkage disequilibrium with them, localise to putative *ZKSCAN3* enhancers that bear active histone marks in human neural progenitors (Figure S16). These findings point to a role of ZKSCAN3 as a neurodevelopmental regulator.

Consistent with this notion, our data demonstrate that ZKSCAN3 is an active enhancer-regulator during neural differentiation. In ZKSCAN3-degraded neural cells, we observed an increased enrichment of H3K4me1 and H3K27ac at ZKSCAN3 binding sites (Figure 5B). Loss of ZKSCAN3-mediated enhancer restriction resulted in neural cells exhibiting a more endodermal identity relative to untreated cells, accompanied by up-regulation of TFs *Gata4*, *Gata6*, *Sox17 and Gdf3.* GDF3, which was directly bound by ZKSCAN3, is a member of the Transforming Growth Factor ² (TGF²) superfamily of cytokines that is necessary for formation of mesoderm and endoderm, as evidenced by the finding that *Gdf3*-null zebrafish embryos completely fail to develop mesoderm or endoderm and consist solely of ectoderm^94^. Expression of endodermal TFs was associated with a deficit in the expression of genes relating to neural processes, including axon development. One such example is *Shh*, a concentration-dependent morphogen and mitogen for which the dose and duration of exposure determines the differential specification of neuronal fates during differentiation^135^. Further evidence that ZKSCAN3-mediated enhancer regulation plays a role in neural differentiation is provided by our observations in pluripotent stem cells undergoing multilineage differentiation. In the neuroectodermal lineage we again observed *Gdf3* upregulation in ZKSCAN3-degraded cells, and this was accompanied by changes to genes related to axonal pathfinding. The Netrin receptors DCC and UNC5 direct the developing axon during neural development, with DCC mediating axon attraction to Netrin and UNC5 mediating repulsion ^102–104,106,107,136,137^. *Dcc* was the most downregulated gene in the neuroectodermal cluster of our scRNA-seq dataset, while *Unc5b* was concurrently upregulated (Figure 6G). By regulating the availability of DCC and UNC5 receptors on the cell surface, the attractive and repulsive actions of Netrin-1 can be modulated^138–140^. Subtle alterations to this ratio during neurodevelopment can result in remodelling of specific neural circuits, with resultant changes in the function of these systems in adulthood^141^. Indeed, mice bred for *Dcc* haploinsufficiency (+/-) exhibit altered development of mesocorticolimbic dopamine neurons and changes to dopamine-related behaviours in adulthood^142,143^, while variations in *Dcc* expression are linked to psychiatric disorders in the human population, including schizophrenia^144^. Thus, during neural differentiation ZKSCAN3 regulates the expression of genes related to axonal development and guidance. This observation suggests a mechanistic basis for the observed genetic association between ZKSCAN3 and schizophrenia in human populations.

In conclusion, our work identifies ZKSCAN3 as a KAP1-independent KZFP that regulates enhancers by targeting embedded SINE elements. Additionally, ZKSCAN3-mediated enhancer regulation during neural specification may suggest a mechanism underlying its genetic association with schizophrenia.

## Acknowledgments

We would like to thank Dr. Helen Rowe for providing the expression plasmid for ZFP809 and for useful discussions. We are grateful to C. D’Santos and V. Roamio Franklin at the Cancer Research UK Cambridge Institute for advice and execution of RIME analyses. We thank B. Hendrich, P. Rugg-Gunn, S. Elderkin, C. Belton and S. Khan for critical review of the work and O. Reos for help throughout the planning and writing of the manuscript. We thank Ciaran Ambler for advice regarding the evolutionary origin of SINE retrotransposons. This work was supported by the Babraham Institute, which receives its core funding from the UK Biotechnology and Biological Sciences Research Council (BBS/E/B/000C0421) and by a Wellcome Trust/Royal Society Sir Henry Dale Fellowship to M.A.C. (105642/Z/14/Z). D.M. was funded by a University of Cambridge MRC DTP studentship. M.S. is supported by UK Medical Research Council Investigator funding (MC-A652-5QA20).

## Author Contributions

D.M. and M.A.C. conceptualized and planned the study. D.M. designed and conducted the experiments. E.W. and S.S. provided technical assistance for the generation of ATAC-seq libraries and advice on the work. S.A. conducted data analysis for scRNA-seq experiments. C.A. assisted in computational analyses. M.S. carried out eQTL mapping and annotation of *Zkscan3* enhancer sites. M.A.C. supervised the work. D.M. and M.A.C. wrote the manuscript, with comments from all authors.

## Data Availability

All sequencing and proteomic datasets associated with this study are deposited in public repositories and will become available upon publication of the manuscript. Requests for access prior to publication can be made to the corresponding author.

## Comparisons with published datasets

Data for expression of *Zkscan3* in the early mouse embryo was sourced from Table S1 of Boroviak et al^32^. P300 ChIP-seq (GSE56098) was sourced from Buecker et al^145^ and ESRRB, OCT4, NANOG and SOX2 ChIP-seq (GSE152186) were sourced form Festuccia et al^61^.

## Supplementary Information

**Table S1** Full RNA-seq quantitation for Zkscan3-*KO* mESCs

**Table S2** Full results from RIME screen for ZKSCAN3 in mESCs

**Table S3** Intersection of significant RIME hits with sysNDD database of neurodevelopmental diseases

**Table S4** Full gene ontology results of up and downregulated genes in ZKSCAN3-degraded neural cells

**Table S5** Full RNA-seq quantitation for ZKSCAN3-degraded neural cells

**Table S6** Primers used in this study

## Materials and Methods

### Cell Culture

All cell types were cultured at 37°C with 5% CO2. E14 mESCs were sourced from the Cambridge Stem Cell Institute, UK. Cells were cultured on 0.1% (w/v) gelatin-coated dishes and passaged every other day or when they reached ∼80% confluency, using TrpLE express enzyme (Gibco, #12604013) to detach. For 2i/LIF conditions, mESCs were maintained in a 1:1 mix of Neurobasal (Gibco, #21103) and DMEM:F12 media (Gibco, 21131), with N2 supplement (Thermo, 17502-048), B27 supplement (Thermo, 17504-044), 2mM L-Glutamine (Gibco, #A2916801), 50μM β-mercaptoethanol (Gibco, #31350010), 0.1mM non-essential amino acids (Gibco, #11140050), 1mM sodium pyruvate (Gibco, #A2916801), 1μM PD0325901 (Sigma, PZ0162), 3μM CHIR99021 (Sigma, SML1046) and 10ng/ml LIF (Cambridge Stem Cell Institute). For Serum/LIF conditions, cells were cultured in Glasgow’s Minimum Essential Medium (GMEM) (Gibco, #11710035) supplemented with 10% Fetal Bovine Serum (Thermo-Fisher), 0.1mM non-essential amino acids (Gibco, #11140050), 1mM sodium pyruvate (Gibco, #11360070), 2mM L-glutamine (Gibco, #A2916801), 100μM β-mercaptoethanol (Gibco, #31350010) and 10ng/ml LIF (Cambridge Stem Cell Institute).

mESCs were differentiated towards the neural lineage by culture in plain N2B27 for six days. N2B27 consisted of a 1:1 mix of Neurobasal (Gibco, #21103) and DMEM:F12 media (Gibco, 21131), with N2 supplement (Thermo, 17502-048), B27 supplement (Thermo, 17504-044), 2mM L-Glutamine (Gibco, #A2916801), 50μM β-mercaptoethanol (Gibco, #31350010), 0.1mM non-essential amino acids (Gibco, #11140050) and 1mM sodium pyruvate (Gibco, #A2916801).

mESCs were differentiated to EBs using the hanging drop method. Cells were resuspended in EB formation medium, which consisted of Glasgow’s Minimum Essential Medium (GMEM) (Gibco, #11710035) supplemented with 10% Fetal Bovine Serum (Thermo-Fisher), 0.1mM non-essential amino acids (Gibco, #11140050), 1mM sodium pyruvate (Gibco, #11360070), 2mM L-glutamine (Gibco, #A2916801), 100μM β-mercaptoethanol (Gibco, #31350010), and plated as hanging drops containing 1000 cells each onto the roof of a bacterial culture dish filled with 5ml of PBS. After two days, the hanging drops were flushed into solution with EB formation medium into fresh bacterial culture plates. Medium was changed after two further days of culture, and EBs were harvested after another two days for a total of six days.

### Generation of *Zkscan3-Fkbpv-HA* cell lines

Cell lines were generated using the protocol described by Arai and Nakao^146^ for efficient biallelic knock-in to mESCs (Figure S2). Briefly, on the first day, cells were reverse transfected with 6μg of plasmid encoding an shRNA for Polq using lipofectamine 2000 reagent (Invitrogen,11668027). On the second day, cells were transfected with 3μg of donor plasmid and 7μg of plasmid encoding Cas9 and an sgRNA, again using lipofectamine. The sgRNA used targeted a site near the *Zkscan3* C terminus (AGTCATTAAGAACTACAGAG). After four hours, the medium was changed to fresh medium with 1μg/mL puromycin and 10μM NU7441 (Cayman Chemical, 14881). On day three and four, medium was changed to fresh medium containing 10μM NU7441, and cells then maintained for a further 6 days in plain medium before FACS sorting single mCherry+ cells into wells of a 96 well plate. The resulting cell populations were screened for knock-in using genomic PCR. Correct integration was confirmed by Sanger Sequencing. To remove the mCherry selection cassette from correct integrants, 2μg of a plasmid encoding Cre recombinase was transfected into 1x10^6^ cells using lipofectamine 2000 and the cells maintained for two days before FACS sorting single mCherry-negative cells into wells of a 96 well plate. The resulting populations were screened by PCR for excision of the selection cassette and confirmed by Sanger Sequencing.

### Generation of *Zkscan3-KO* lines

sgRNA sequences (see table below) were ordered as gBlock constructs according to the design below. Targeting sequence (green) was included downstream of a U6 promoter (yellow) and upstream of sgRNA scaffold sequence (blue) and terminator sequence (red).

TGTACAAAAAAGCAGGCTTTAAAGGAACCAATTCAGTCGACTGGATCCGGTACCAAGGTCGGGCAGGAAGAGGGCCTATTTCCCATGATTCCTTCATATTTGCATATACGATACAAGGCTGTTAGAGAGATAATTAGAATTAATTTGACTGTAAACACAAAGATATTAGTACAAAATACGTGACGTAGAAAGTAATAATTTCTTGGGTAGTTTGCAGTTTTAAAATTATGTTTTAAAATGGACTATCATATGCTTACCGTAACTTGAAAGTATTTCGATTTCTTGGCTTTATATATCTTG TGGAAAGGACGAAACACC**XXXXXXXXXXXXXXXXXXX**GTTTTAGAGCTAGAAATAGCAA GTTAAAATAAGGCTAGTCCGTTATCAACTTGAAAAAGTGGCACCGAGTCGGTGCTTTTTTTCTAGACCCAGCTTTCTTGTACAAAGTTGGCATTA

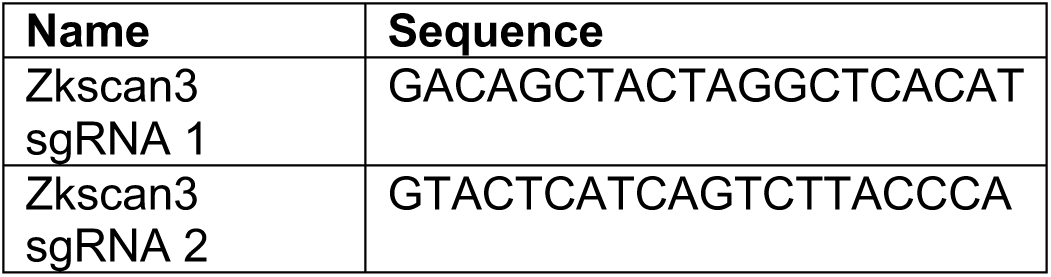

1x10^6^ WT E14 mESCs were transfected with 600ng hCas9-GFP plasmid and 100ng of gBlock with Lipofectamine 2000 (Invitrogen,11668027). Medium was changed to fresh medium after 6hr. After 48hr of culture, the cells were sorted for GFP+ cells using Fluorescence Activated Cell Sorting (FACS) into 96 well plates as single cells. Colonies were then screened for knockout by flanking genomic PCR followed by Sanger Sequencing to check for indels. Primer sequences used for PCR are detailed in Table S6. Sanger Sequencing was performed in both directions, using the same primers as the flanking PCR, by Azenta Sequencing Services UK.

### Immunoblotting

Whole cell extracts were generated from cells washed once in ice-cold Phosphate Buffered Saline (PBS) followed by direct addition of 2X Laemmli buffer (120mM Tris-HCl pH 6.8, 4% (w/v) SDS, 20% (v/v) glycerol) to the cell culture vessel. Extracts were scraped into ice-cold 1.5ml Eppendorf tubes, boiled for 5 minutes and homogenized with a 25G needle. Protein concentrations were quantified using a NanoDrop 2000 spectrophotometer. Samples were either analyzed immediately or stored at -80°C.

Protein concentration was equalised with water and 2x Laemmli Sample Buffer (65.8 mM Tris-HCl pH 6.8, 26.3% (w/v) glycerol, 2.1% (w/v) SDS, 0.01% (w/v) bromophenol blue; Biorad #1610737) supplemented with β-mercaptoethanol to a final concentration of 355 mM. 30μg of protein was loaded into each well of a 4-20% TGX Criterion gradient gel (BioRad, #5671094). Samples were then resolved on the gel at 120V for 1-1.5 hours with Tris-Glycine SDS Running buffer (25mM Tris-HCl, 190mM glycine, 0.1% (w/v) SDS). Proteins were transferred onto 0.2μm pore nitrocellulose membranes for one hour at 800mA with Tris-Glycine Transfer buffer (25mM Tris-HCl, 190mM glycine, 20% (v/v) methanol). Transfer efficiency was assessed by staining the membrane with Ponceau S stain (0.1% Ponceau S (w/v), 5% acetic acid (v/v)) which was washed away with TBS-T (Tris-buffered saline, 0.1% (v/v) TWEEN-20) under gentle agitation on a rocking platform. Membranes were incubated with blocking buffer (5% (w/v) Bovine Serum Albumin (BSA) in TBS-T) for one hour at room temperature on a rocking platform. Primary antibodies were added at the appropriate concentration (see table below) in Blocking Buffer and incubated on a rocking platform in a 4°C cold room overnight. The next day, membranes were washed 3 times for 5 minutes in TBS-T and HRP conjugated secondary antibody added at the appropriate concentration, diluted in blocking buffer, for one hour at room temperature on a rocking platform. Membranes were again washed 3 times for 5 minutes in TBS-T then incubated with SuperSignal™ West Pico PLUS Chemiluminescent Substrate (Thermo, #34579) before being imaged on a G:BOX Chemi visualiser (Syngene).

| Antibody | Supplier/Catalogue number | Dilution |
| --- | --- | --- |
| H3K9me3 | Abcam (ab176916) | 1:5000 |
| HA tag | Cell Signalling Technology (3724) | 1:1000 |
| KAP1 (TRIM28) | Thermo (PA527648) | 1:1000 |
| GAPDH | Cell Signalling Technology (2118) | 1:5000 |
| Anti-rabbit IgG HRP | Abcam (Ab6721) | 1:3000 |
| Anti-mouse IgG HRP | Abcam (Ab6728) | 1:2000 |

### Reverse Transcription Quantitative Polymerase Chain Reaction (RT-qPCR)

RNA was extractedfrom cells using the Qiagen RNeasy Mini Kit (Qiagen, # 74104). RNA concentrations were quantified with a NanoDrop 2000 spectrophotometer. To generate cDNA, 1μg of extracted RNA was processed with the QuantiTect Reverse Transcription Kit (Qiagen, #205311). Remaining RNA was stored at -80°C. cDNA was quantified with a NanoDrop 2000 spectrophotometer and aliquots diluted to 20ng/μl in preparation for qPCR. Remaining cDNA was stored at -20°C.

Primers for each target (see Table S6) were diluted into master mixes with nuclease-free water and SYBR^®^ Green JumpStart *Taq* ReadyMix (Sigma, #S4438). Primers were used at a final concentration of 200nM. 7μl of master mix and 5μl of cDNA, for a total of 100ng cDNA, was added to each well of a 96 or 384 well plate with an electronic pipette for a total volume of 12μl per reaction. Samples were processed as both biological and technical triplicates. Samples were mixed with a multichannel pipette, placed in a plate centrifuge for 1 minute at 1000xg, and the plate sealed before being placed into a BioRad CFX96/384 Real Time PCR Detection System with the following programme: 94°C for 2 minutes, followed by 40 cycles of: 94°C for 15 seconds, 55°C for 1 minute. Melt curve analysis was performed after the run to check for the presence of a single amplicon. No-Template controls were run alongside samples to check for extraneous nucleic acid contamination or primer-dimer formation. Relative gene expression values were calculated using the Pfaffl method^147^. Expression values were normalised to housekeeping genes as indicated in figure legends.

### RNA-sequencing

RNA was extracted from cells using the Qiagen RNeasy Mini Kit (Qiagen, 74104). RNA concentrations were quantified with a NanoDrop 2000 spectrophotometer. For RNA-seq from WT and *Zkscan3-KO* mESCs, libraries were prepared using the NEBNext rRNA Depletion Kit (New England Biolabs, E6310) and NEBNext Ultra II Directional RNA Library Prep Kit for Illumina (New England Biolabs, E7760). Libraries were quantified using the KAPA Library Quantification Kit (KK4873) according to manufacturer’s instructions and diluted to 5nM for sequencing (performed by the CRUK CI Genomics facility) as 150bp PE reads on a NovaSeqX instrument.

For RNA-seq of *Zkscan3-Fkbpv-HA* cells upon neural differentiation, the NEBnext PolyA mRNA magnetic isolation module was used, followed by input to the NEBNext Ultra II Directional RNA Library Prep Kit for Illumina (New England Biolabs, E7760). Libraries were quantified using the KAPA Library Quantification Kit (KK4873) according to manufacturer’s instructions and diluted to 5nM for sequencing (performed by the Babraham Institute Genomics Facility) as 150bp PE reads on an AVITI sequencing instrument.

Quality and adapter trimming were performed withTrim Galore v0.6.5 (using Cutadapt v2.3). Mapping to the mouse reference genome (GRCm39) was performed using Hisat2 v2.2.1 ^148^. Reads were quantified using the RNA-seq quantitation pipeline in the SeqMonk tool for opposing strand specific libraries and paired-end reads. Differential expression was assessed using DESeq2^149^ with an adjusted p value cutoff of 0.05.

### ChIP-sequencing

Cells were fixed for 8 minutes with 1% formaldehyde (methanol free) (Thermo, 11586711) and the reaction quenched with 125mM glycine for 5 minutes. Cells were washed twice with PBS and scraped in 10ml of PBS into falcon tubes, and cell pellets collected by centrifugation at 300xg for 5 minutes at 4°C. Pellets were suspended in Lysis Buffer 1 (10mM Tris-CL pH 8.0, 10mM EDTA, 0.25% Triton X-100) and incubated on ice for 10 minutes. The buffer was supplemented with Protease Inhibitor Cocktail (PIC) (Sigma, P8340). Nuclei were pelleted by centrifugation at 500xg for 5 minutes at 4°C. Nuclei were resuspended in Lysis Buffer 2 (10mM Tris-Cl pH 8.0, 10mM EDTA, 0.25% Triton X-100, 0.1% SDS, 200mM NaCl, 1X PIC) and incubated on ice for 10 minutes. Cells were distributed into 1.5ml Bioruptor TPX tubes (Diagenode, C30010010) such that ∼10 million cells were present in each tube in 300μl volume. Chromatin was sheared using a Diagenode Bioruptor Plus instrument for 2x10 cycles of 30 seconds ON / 30 seconds OFF on the HIGH power setting. Debris was cleared by centrifugation at maximum speed in a benchtop centrifuge for 10 minutes. Chromatin shearing was assessed each time the procedure was performed to ensure proper fragment sizes were obtained. Meanwhile, chromatin was frozen at -80°C. To check shearing, an aliquot was removed from the sheared chromatin, diluted in SDS-containing buffer (100 mM Tris pH8, 1%SDS, 200 mM NaCl) and incubated with 0.8 units molecular-biology grade proteinase K (New England Biolabs, P8107S) overnight at 65°C on a thermal shaker to reverse crosslinks. The following day, 20μg of RNAse A (New England Biolabs, T3018-2) was added to each sample and incubated for one hour at 37°C on a thermal shaker. A further 0.8 units of molecular-biology grade proteinase K was added and the sample incubated for a further 30 minutes on a thermal shaker at 55°C. DNA was cleaned up using the QiaQuick PCR purification kit (Qiagen, 28104) and the size distribution of fragments checked by agarose gel electrophoresis. If shearing was successful then chromatin stocks were defrosted on ice and the volume of each sample made up to 1ml with Dilution Buffer (20mM Tris-Cl pH 8.0, 2mM EDTA, 150mM NaCl, 1X PIC). DynaBeads Protein G (Invitrogen, 10003D) were washed once with Dilution Buffer and then 25μl of beads were added to each sample for one hour on a rotator at 4°C, to pre-clear the suspension. Beads were removed and discarded using a magnetic rack, and antibodies were added overnight to immunoprecipitate targets (see table below). Samples were rotated at 4°C overnight. Meanwhile, beads were blocked overnight in 7.5% BSA in PBS. The next day, blocked beads were resuspended in Dilution Buffer and 25μl added to each sample for four hours on a rotator at 4°C. Beads were then washed sequentially with Wash Buffer 1 (20mM Tris-Cl pH 8.0, 2mM EDTA, 150mM NaCl, 1% Triton, 0.1% SDS, 1X PIC), Wash Buffer 2 (20mM Tris-Cl pH 8.0, 2mM EDTA, 300mM NaCl, 1% Triton, 0.1% SDS, 1X PIC) and Wash Buffer 3 10mM Tris-Cl pH 8.0, 2mM EDTA, 250mM LiCl 0.5% NP-40, 0.5% deoxycholate, 1X PIC) with each wash lasting 15 minutes at 4°C on a rotator. Beads were washed once in TE buffer (10mM Tris-Cl, pH 8.0, 1mM EDTA). DNA was eluted by adding 100ul Elution Buffer (1% SDS, 0.75% sodium bicarbonate) supplemented with 0.4 units proteinase K to each sample and incubating for 15 minutes at 55°C on a thermal shaker. This eluate was removed and a second elution performed with another 100μl Elution Buffer and 0.4 units proteinase K. The two eluates were pooled and incubated overnight at 65°C on a thermal shaker. The following day, 20ug of RNAse A was added to each sample and incubated for one hour at 37°C on a thermal shaker. A further 0.8 units of molecular-biology grade proteinase K was added and the sample incubated for a further 30 minutes on a thermal shaker at 55°C. DNA was cleaned up using the QiaQuick PCR purification kit and quantified using the QuBit hsDNA high sensitivity kit (Invitrogen, Q33230). Libraries were prepared using the NEB Next Ultra II Library Prep Kit for Illumina (New England Biolabs, E7645), quantified using the KAPA Library Quantification Kit (KK4873) according to manufacturer’s instructions and diluted to 5nM for sequencing (performed by the Babraham Institute Genomics Facility) as 150bp PE reads on an Illumina HiSeq instrument (mESC ChIPs) or AVITI sequencing instrument (all neural differentiation ChIPs).

### ATAC-seq

ATAC-seq was performed using the “Omni-ATAC” protocol from Corces and colleagues ^150^. To begin, 50,000 cells were resuspended in 1ml of cold Resuspension Buffer (10mM Tris-HCl pH 7.4, 10mM NaCl, 3mM MgCl_2_) and then centrifuged at 500xg for 5 minutes at 4°C. The cell pellets were then resuspended in 50μl of Resuspension Buffer with the addition of 0.1% NP40, 0.1% Tween-20, and 0.01% digitonin and incubated on ice for 3 min. Next, 1ml of cold Resuspension Buffer containing 0.1% Tween-20 (without digitonin or Tween) was added and the tube was inverted to mix. Nuclei were pelleted by centrifugation for at 500xg for 10 minutes at 4°C. The nuclei were then resuspended in 50μl of transposition mix (25μl 2x TD buffer [20mM Tris-Cl pH 7.6, 10mM MgCl2, 20% dimethyl formamide], 2.5μl transposase (100nM final), 16.5ul PBS, 0.5μl 1% digitonin, 0.5μl 10% Tween-20, 5μl H2O). This suspension was incubated for 30 minutes at 37°C on a thermal shaker. Samples were cleaned using Zymo DNA Clean and Concentrator-5 columns (Zymo Research, D4014). To the ∼20μl of product, 2.5μl of 25μM i5 primer and 2.5μl of 25μM i7 primer and 25μl of 2X NEBNext high fidelity master mix (NEB, M0541) was added. 5 cycles of PCR were performed with conditions: 72°C for 5 minutes; 98°C for 30 seconds; then 5 cycles of 98°C for 10 seconds, 63°C for 30 seconds and 72°C for 60 seconds. Next a qPCR reaction was performed to determine the additional number of amplification cycles necessary. 5μl (∼10%) of the pre-amplified sample was combined with 3.76μl of water, 0.5μl 25μM i5 primer, 0.5μl 25μM i7 primer, 0.24μl of 25X SYBR Green (ABP Biosciences, D010) and 5μl 2X NEB master mix and qPCR performed with conditions: 98°C for 30 seconds followed by 20 cycles of [98 °C for 10 seconds, 63°C for 30 seconds, 72°C for 60 seconds]. To determine the additional number of cycles, linear fluorescence was plotted against cycle number, and the cycle number that corresponded to 1/3 of the maximum fluorescence intensity was chosen as the number of extra cycles. The remainder of the pre-amplified DNA was then run in the thermocycler for the additional number of cycles, cleaned up again with Zymo DNA Clean and Concentrator-5 columns and a single 1.2X cleanup was performed using AMPure beads. Libraries were quantified using the KAPA Library Quantification Kit (KK4873) according to manufacturer’s instructions and diluted to 5nM for sequencing (performed by the Babraham Institute Genomics Facility) as 150bp PE reads on an Illumina HiSeq instrument.

### RIME

RIME was performed according to the protocol developed by D’Santos and colleagues ^70^. Briefly, Protein G dynabeads (Invitrogen, 10003D) were washed 4 times in PBS supplemented with 5mg/ml BSA, resuspended in the same PBS/BSA solution and 10μg of primary antibodies (anti-HA [Ab9110] or anti-IgG [Ab172730]) added overnight at 4°C on a rotator. The next day, all buffers were filtered and protease inhibitor cocktail (PIC) was added (Sigma, P8340), and they were pre-chilled to 4°C. Cells were washed in PBS and then fixed for 8 minutes with 1% formaldehyde (methanol free) (Thermo, 11586711) and the reaction quenched with 125mM glycine for 5 minutes. Cells were washed twice with 10ml of ice-cold PBS. Using a silicone scraper, cells were collected in 10ml of PBS and pelleted by centrifugation at 2,000g for 3 minutes at 4°C in a 15-ml conical tube. The supernatant was removed, and the cells were resuspended in 10ml of PBS. Centrifugation was repeated, and the final PBS wash was discarded. The cell pellet was resuspended in 100ml of Lysis Buffer 1 (50mM HEPES-KOH, (pH 7.5), 140mM NaCl, 1mM EDTA, 10% (vol/vol) glycerol, 0.5% (vol/vol) NP-40/Igepal CA-630, 0.25% (vol/vol) Triton X-100, 1X PIC) and rotated 10 minutes on a rotator at 4°C. The lysate was cleared by centrifugation at 2,000g for 5 minutes at 4 °C. The pellet was resuspended in 10ml of Lysis Buffer 2 (10mM Tris-HCL (pH 8.0), 200mM NaCl, 1 mM EDTA and 0.5mM EGTA) and rotated 5 minutes at 4°C. The lysate was cleared by centrifugation at 2,000xg for 5 minutes at 4°C. An equivalent of 1x10^7^ cells was resuspended in 300μl of Lysis Buffer 3 (10mM Tris-HCl (pH 8.0), 100mM NaCl, 1mM EDTA, 0.5mM EGTA, 0.1% (wt/vol) sodium deoxycholate and 0.5% (vol/vol) *N*-lauroylsarcosine, 1X PIC) in 1.5ml Bioruptor TPX tubes (Diagenode, C30010010). The suspension was sonicated using a Diagenode Bioruptor Plus instrument for 10 cycles of 30 seconds ON / 30 seconds OFF on the HIGH power setting. To the sonicated lysate, 30μl of 10% (vol/vol) Triton X-100 was added, vortexed, and the lysate clarified by centrifugation at 20,000xg for 10 minutes at 4°C. The supernatant was transferred to a fresh tube. The antibody-bound beads were retrieved from the overnight incubation washed in 1ml of PBS/BSA five times before being resuspended in 100μl of PBS/BSA which were to each sample and rotated at 4°C overnight. The following day, the beads were washed in 1ml of RIPA buffer (50mM HEPES (pH 7.6), 1mM EDTA, 0.7% (wt/vol) sodium deoxycholate, 1% (vol/vol) NP-40, 0.5M LiCl, 1X PIC) ten times. The beads were then washed twice in 1ml of freshly made, cold 100mM ammonium hydrogen carbonate (AMBIC) solution, and the beads transferred to fresh tubes after the second wash, frozen on dry ice and transferred to the Mass Spec Core at CRUK Cambridge Institute, UK.

To digest the bead-bound protein, 100ng of trypsin (Pierce, #90058) in 100mM ammonium bicarbonate (AMBIC) was directly added to the AMBIC-washed beads The samples were vortexed for 15 seconds every 2–3 minutes for the first 15 minutes to ensure even suspension of the beads in the trypsin solution and then left to digest overnight at 37°C. Following the overnight digestion, an additional 100ng of trypsin was added to each sample and the digestion was continued for another 4 hours at 37°C. The tubes were then placed back on the magnetic rack to remove the supernatant, which was subsequently added to a tube containing 5% trifluoroacetic acid (TFA) to achieve a final TFA concentration of 0.5% (vol/vol) to stop the reaction.

For desalting the digested samples, C18 cartridges (Pierce, #84850) were conditioned with 2 × 20μl of 80% (vol/vol) acetonitrile/0.1% TFA. The cartridges were then equilibrated with 2 × 20μl of 0.1% (vol/vol) TFA. The acidified peptides were loaded onto the cartridges, which were subsequently washed three times with 0.1% (vol/vol) TFA. Peptides were eluted three times with 20μl of 80% (vol/vol) acetonitrile/0.1% (vol/vol) TFA. The eluates were combined and the peptides were dried using a centrifugal vacuum concentrator.

The peptide samples were reconstituted in 25μl of 0.1% (vol/vol) formic acid, and 5μl was injected onto a Dionex Ultimate 3000 UHPLC system coupled with a nano-electrospray ionization (nano-ESI) Fusion Lumos (Thermo Scientific) mass spectrometer. Peptides were loaded and separated on a reverse-phase trap column (2cm, 100 μm i.d.) and an analytical column (25cm × 75μm i.d., BEH130, Waters) using a 5–45% acetonitrile gradient in 0.1% formic acid at a flow rate of 300nl/min. In each data collection cycle, one full MS scan (400– 1,600m/z) was acquired in the Orbitrap at a resolution of 60,000 with an automatic gain control (AGC) setting of 3×10^6^ and a maximum injection time (MIT) of 100ms. Data-independent acquisition (DIA)-MS2 was performed in the range of m/z 380-980 using an isolation window of 10 m/z for 60 scan events with a collision energy of 32%, an AGC setting of 200%, and an MIT of 40ms with loop control of 30 spectra.

Raw data processing was conducted using the DIA-NN software in library-free mode, utilizing the Mus musculus fasta file (taxon ID 10090 -Version October 2023). Trypsin was set as the enzyme with allowance for 2 missed cleavages, and oxidation of methionine was specified as a dynamic modification. The false discovery rate (FDR) cut-off was set at 0.01 at the precursor level. Match between runs (MBR) was enabled for a library-free quantitative analysis to ensure stringent FDR control. Additional options selected included "process in batches," "use neural network," "use isotopologues," "RT profiling," and "remove likely interference." Peptide precursors (pr.matrix) and protein matrices (pg.matrix) were used as the result files. The mean intensity of a peptide’s precursors with an identical stripped sequence was used as the peptide’s intensity for further analysis.

To assess significant interactors, the protein matrices (pg.matrix) results file was input to the DEP2 package^151^. Proteins were filtered for those present in at least 4 replicates. Data were normalised using the normalise_vsn command and missing values imputed using “MinDet”. Significant interactors were defined as those with an adjusted p value of <0.05 and a log2 fold change >1.

### scRNA-seq

A P100 wide-bore pipette tip was used to collect uniformly sized EBs (avoiding any large aggregates) into a 1.5ml DNA LoBind tube (Eppendorf). The EBs were washed twice in PBS using a wide bore pipette tip to resuspend the cells and centrifugation at 150xg for 3 minutes at 4°C. 1ml of pre-warmed (37°C) TrypLE (Gibco, #12604013) was added to the EBs, and they were placed in a thermal shaker for 10 minutes at 37°C with 1500 rpm shaking. After 10 minutes, a normal P1000 tip was used to pipette up and down five times to break up any remaining cell clumps. The cells were washed twice in PBS/0.04% BSA using a wide bore pipette tip to resuspend the cells and centrifugation at 150xg for 3 minutes at 4°C. The cell suspension was passed through a 30μm Celltrics filter (Sysmex, 04-004-2326), using a normal P1000 tip to force the liquid through the mesh. Cells were counted and the viability assessed using a Countess Cell Counter (Invitrogen). Viability was above 90% for all samples. The cells were re-suspended to ∼1500cellsμl in 40μl PBS/0.04% BSA. A total of 16,520 cells for untreated samples and 17,700 cells for dTAG treated samples were provided as single cell suspensions and underwent library preparation using the 10x Chromium Next GEM Single Cell 3’ kit v3.1 (dual index) for gene expression libraries. Of these cells loaded onto the chip, approximately 8,260 for untreated samples and 8,850 for dTAG-treated samples were recovered from the chip following manufacturers recommendations for Gel Bead in Emulsion (GEM) generation. The remainder of the 10x library preparation was carried out as per the manufacturers protocol. Resulting gene expression libraries were quality checked on the Agilent Bioanalyzer 2100 using the high sensitivity DNA assay and sequenced on a high output flow cell on the AVITI sequencer.

Raw reads were processed using CellRanger v8.0.0 using the 10X GRCm39-2024-A reference genome. Analysis was performed in R using Seurat v5. Samples were filtered to remove cells with a log10 ratio of counts over median counts of above 0.5 or below -0.3. Cells with mitochondrial sequence content of <1% or >6% were also removed. Raw counts were normalised to log10 reads per 10,000 reads. Variable genes were defined using the variance stabilising transform method and selecting the top 2000 most variable genes and then filtering for those with log2 standardised variance of >1, producing a set of 1483 variable genes.

The data was then scaled to z-scores before PCA analysis on the variable features. The top 12 principle components were used for UMAP dimensional reduction using 20 nearest neighbours and a minimum distance of 0.1. Cells were divided into 19 clusters using Louvain clustering with a resolution of 1 using the same 12 PCA dimensions. Per-Cluster markers were defined using a wilcoxen signed rank test, comparing the expression of the genes in one cluster to cells in all other clusters. P-values were corrected using Benjamini and Hochberg false discovery rate correction. Significant markers were deemed to have an adjusted p-value below 0.01.

Cluster marker gene set analysis was performed using GOST from Gprofiler using gene sets from Gene Ontology Biological Process. The background comprised all genes observed in at least 5% of the cells. Multiple testing correction again used false discovery rate transformation. Per-Cluster differential expression between WT and degron samples was similarly performed using the Wilcoxon test. Significant hits were deemed to have an adjusted p-value of <0.05 and and average log2 fold change of >0.5. Per-Cluster differential expression gene set analysis was performed as before, using the same background as the cluster marker analysis.

Clustering of Gene Ontology groups was performed by calculating the maximum percentage overlap of hit genes in each pairwise combination of terms. Selection of transcriptional regulators came from genes annotated to the Transcriptional Regulator Gene Ontology Category in Ensembl v112. The mouse gastrulation data was obtained from the BioConductor MouseGastrulationData package using all deposited samples and using the pre-calculated UMAP projection and clusters.

### ChIP and ATAC-sequencing processing

Quality and adapter trimming were performed using Trim Galore v0.6.5 (using Cutadapt v2.3). Mapping to the mouse reference genome (GRCm39) was performed using Bowtie2 v2.4.1^152^. Peak calling was performed with MACS2 (v2.2.9.1)^153^ with the options *-f BAMPE --broad -g mm* (H3K7Ac, H3K9me3, H3K4me1), *-f BAMPE –broad -g mm –-nomodel* for ATAC seq or *-f BAMPE -g mm* (all other ChIPs). Troublesome regions were filtered with a custom blacklist, based on the relevant input control, using the SeqMonk tool. For calling ZKSCAN3 peaks in mESCs, all four replicates were considered. Peaks were called permissively for each replicate using MACS2 *(-g mm -f BAMPE -p 1e-2*) against the input control. These permissively called peaks were then filtered with the MSPC package^154^, which evaluates the statistical significance of repeated evidence of ChIP-seq peaks across multiple replicates, thus “rescuing” weak peaks and reducing false negatives. MSPC was run with options *-r bio -w 1e-4 -s 1e-8 -c 3*. Further filtering was applied in the SeqMonk tool, to remove regions with exceptionally high input signal or very low enrichment. For neural lineage cells, two replicates were considered and only MACS2 called peaks (*-f BAMPE -g mm*) occurring in both replicates were retained. Motif analysis was performed using the MEME and SEA tools in the MEME suite,^155^ with standard parameters. For SEA, the sequences were searched against the HOCOMOCO Mouse (v11 CORE) database^50^. For calling differential enrichment of ChIP-seq or ATAC-seq reads, the limma package was used with an adjusted p value cutoff of 0.05. For calling differential enrichment of peaks in neural lineage cells, libraries for each modification were enrichment normalised to the 50^th^ and 90^th^ percentile using the SeqMonk tool. H3K27Ac and ATAC rescue libraries were also enrichment normalised against untreated and dTAG treated libraries, to the 50^th^ and 90^th^ percentile. For visualisation, BigWig files were generated using the BamCoverage tool from DeepTools^156^ with the options *--normalizeUsing CPM --minMappingQuality 20*.

### SNP analysis

The locations of GWAS SNPs from Chen et al^31^ were remapped onto the hg38 assembly using dbSNP 155. SNPs in tight linkage disequilibrium with these SNPs (R2>0.9) were identified using LD linkusing^157^ high coverage hg38 data for all populations. Associations with *ZKSCAN3* expression in Brain_Frontal_Cortex_BA9 were confirmed using GTEx online tool^158^. Histone modification ChIP-seq data for the human mid-frontal lobe region from the Roadmap Epigenomics consortium^159^ were downloaded as wiggle tracks from GEO accessions GSM1112810 (H3K27ac) and GSM670015 (H3K4me1), respectively, converted to BED format using BEDOPS wig2bed ^160^and lifted over from hg19 to hg38 using UCSC liftOver (^161^Data were plotted using the R package plotgardener ^162^.

