## Supplementary Figures for "The KRAB-Zinc Finger protein ZKSCAN3 represses enhancers via embedded retrotransposons"

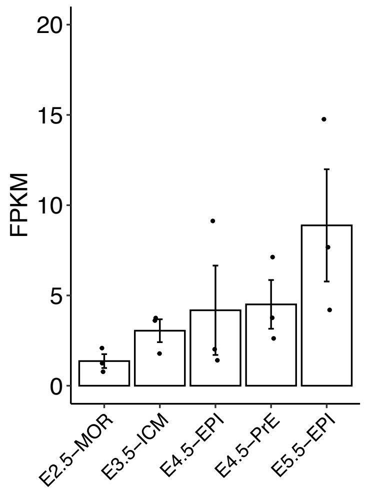


**Figure S1. Expression of Zkscan3 during early mouse embryonic development.**
Expression of *Zkscan3* at early embryonic stages from lineage-specific RNA sequencing[32]. Each point represents a single embryo. Error bars show SEM. Column heights represent the mean. FPKM = Fragments Per Kilobase per Million reads. MOR = Morula, ICM = Inner Cell Mass, EPI = Epiblast, PrE = Primitive Endoderm.

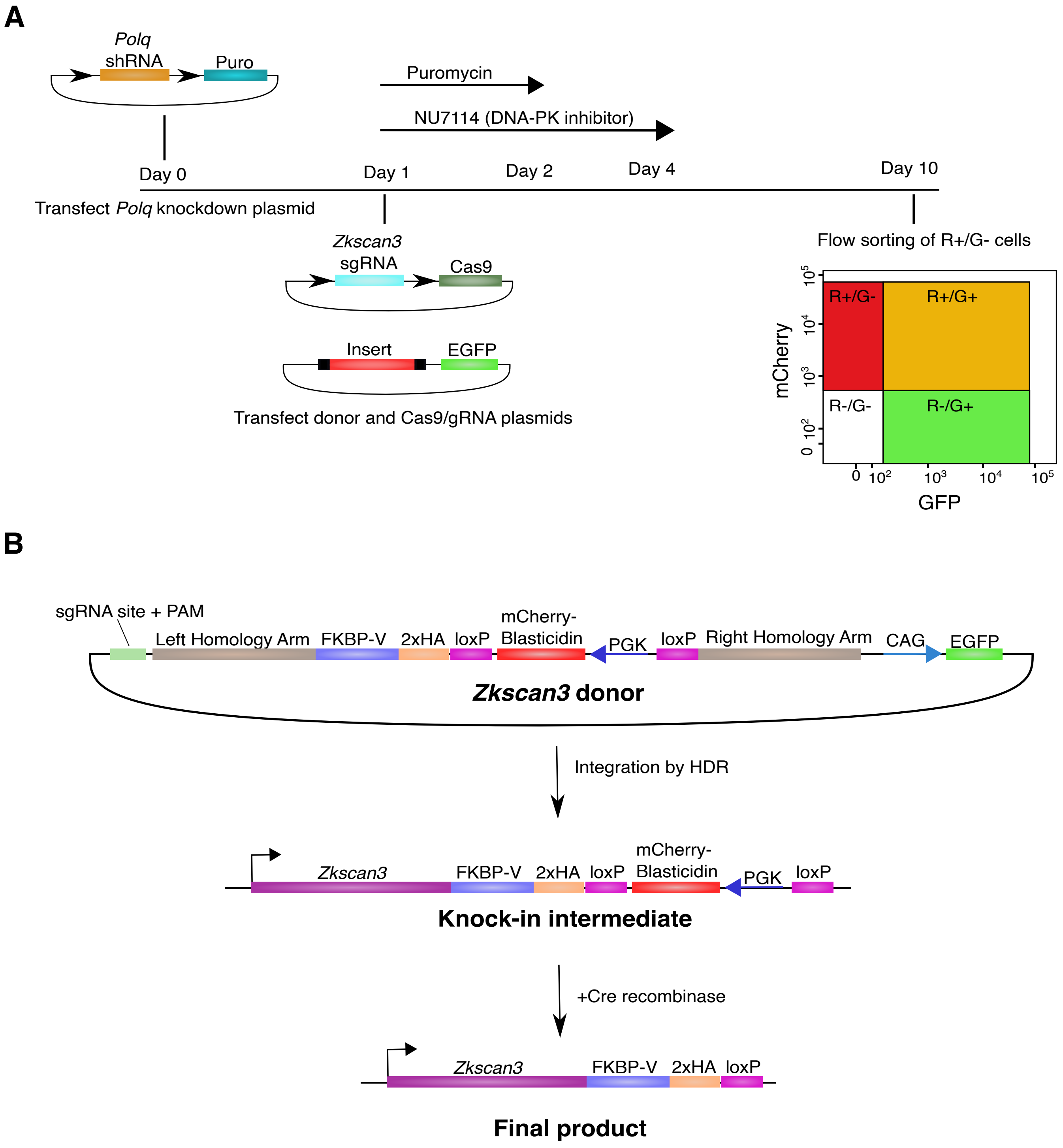


**Figure S2 | Bi-allelic knock-in of FKBPv degron and HA tags to the endogenous *Zkscan3* locus**

**(A)** Schematic outlining the BiPoD process[142], which promotes knock-in via the HDR repair pathway by dual inhibition of Polθ and DNA-PK using shRNA and an inhibitor, respectively. Cells are screened for integration of the donor DNA with flow cytometry, where cells positive for mCherry but negative for GFP, are retained as correct integrants. **(B)** Schematic outlining the design of the donor plasmid for Homology Directed Repair (HDR), the resulting knock-in genotype, and the final genotype after Cre recombianse mediated excision of the mCherry-blasticidin selection casette. An sgRNA cut-site and PAM for the *Zkscan3* sgRNA is included on the donor to facilitate linearisation of the plasmid inside the cell, which enhances knock-in efficiency. GFP, located outside the homology arms, is used to screen out integrations of the whole donor plasmid

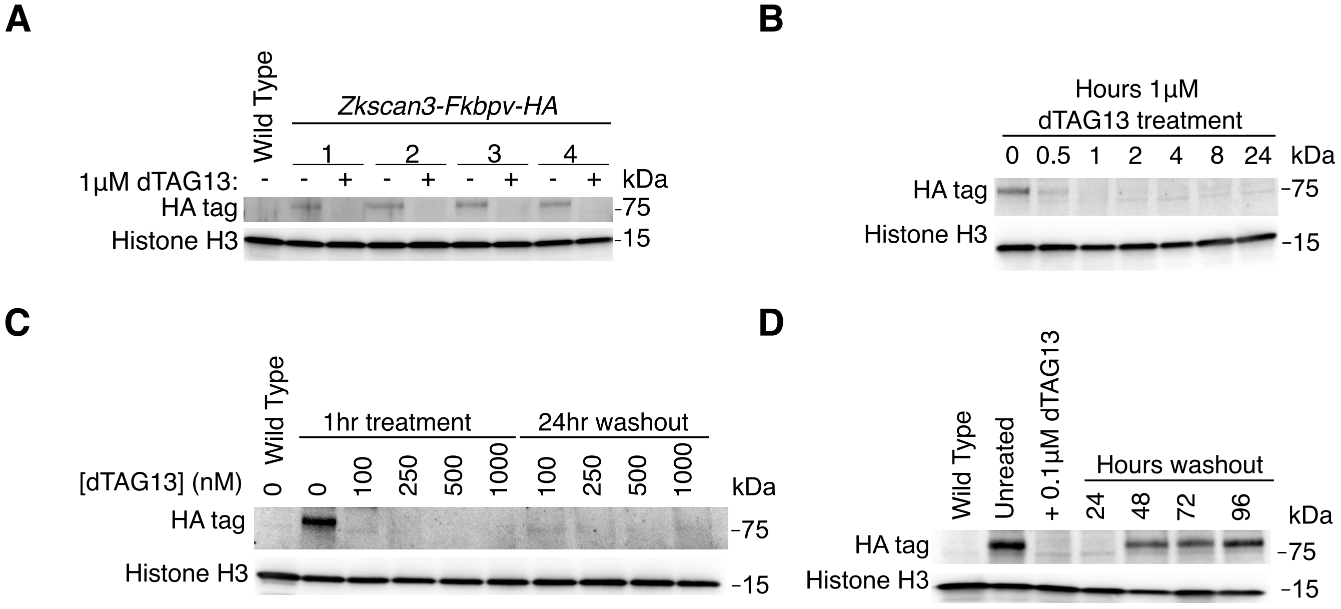


**Figure S3. Rapid degradation and recovery of ZKSCAN3 in *Zkscan3-Fkbpv-HA* knock-in mESCs**

**(A)** Immunoblot showing degradation of ZKSCAN3 following treatment with 1µM dTAG13 for 24 hours in four independent clonal lines. **(B)** Immunoblot showing time-course of ZKSCAN3 degradation in *Zkscan3-Fkbpv-HA* cells treated with 1µM dTAG13. **(C)** Immunoblot showing degradation of ZKSCAN3 in *Zkscan3-Fkbpv-HA* cells treated with different amounts of dTAG13. Cells were treated for 1 hour. 24 hour washout refers to 24 hours after removal of dTAG13 from the culture medium. **(D)** Immunoblot showing re-establishment of ZKSCAN3 expression after removal of dTAG13 from culture medium. *Zkscan3-Fkbpv-HA* cells were treated with 100nM dTAG13 for 1 hour, after which the cells were washed, the medium changed, and the cells allowed to grow for 24, 48, 72 or 96 hours before harvest.

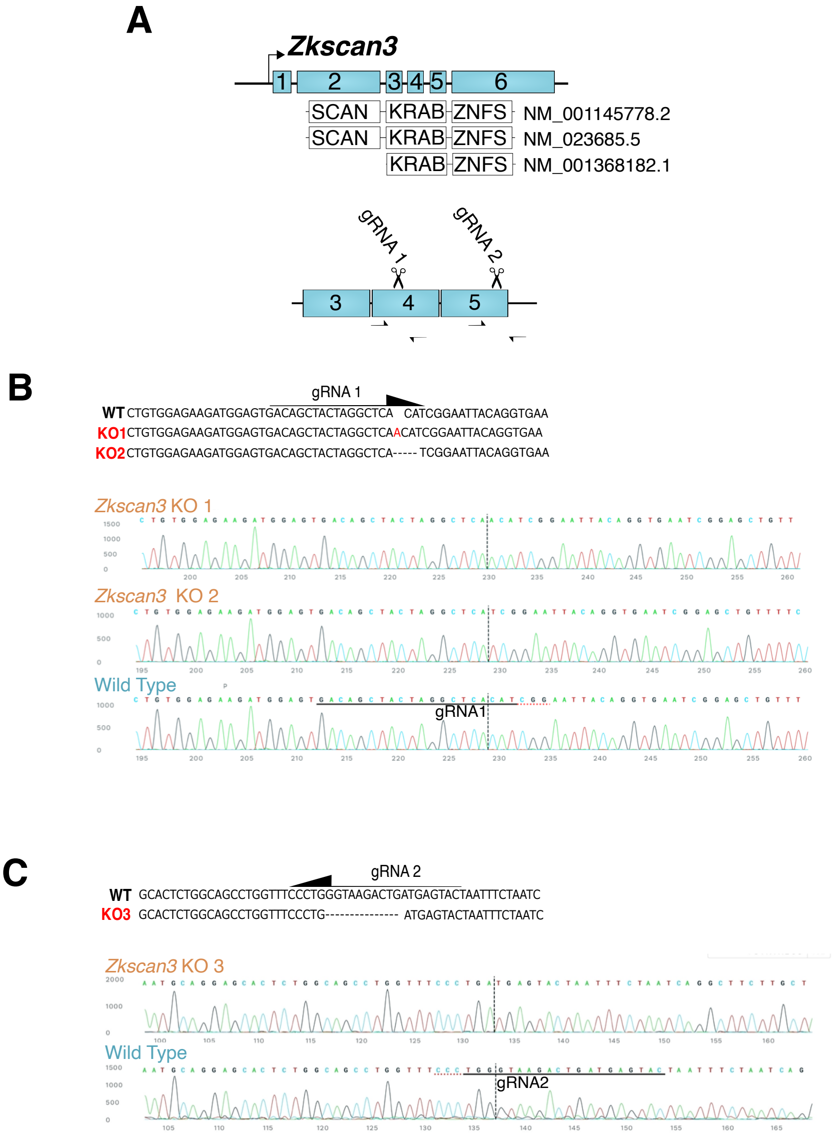


**Figure S4. Generation of *Zkscan3* KO mESCs**

**(A)** Schematic showing the *Zkscan3* gene and curated transcripts, and the location of sgRNAs designed to knock out expression of all curated transcripts. The location of PCR primers used in screening is indicated by arrows. **(B,C)** Results of Sanger sequencing screening for *Zkscan3* knockout with gRNA1 (B) and gRNA2 (C). Top: summary of the genotypes of the knockout clones generated, showing indels relative to the WT sequence. Bottom: Sanger sequencing traces showing the relevant indels.

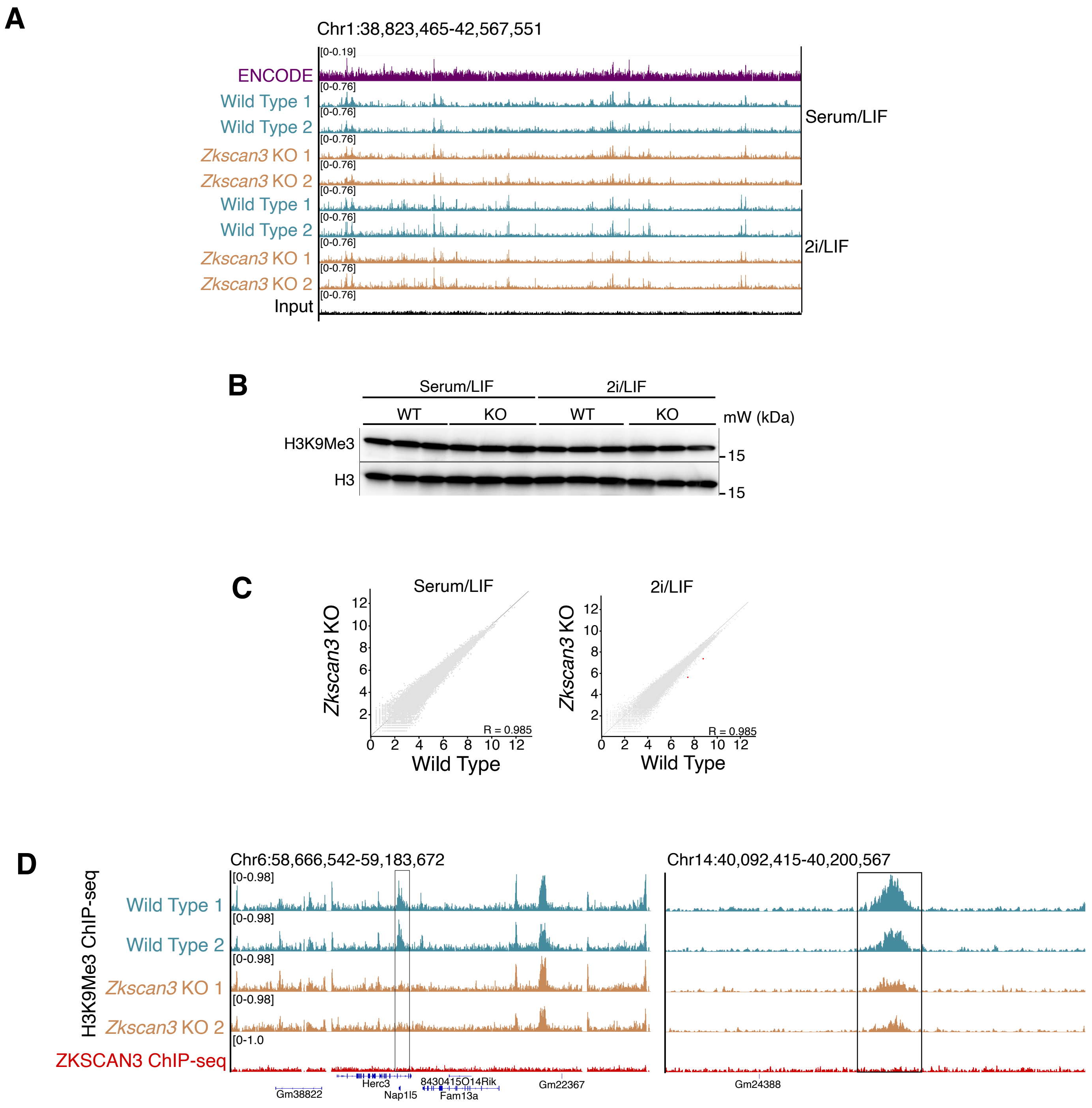


**Figure S5. Zkscan3 knockout does not widely affect H3K9me3 localisation in mESCs**

**(A)** Genome browser tracks showing H3K9me3 ChIP-seq signal for WT and *Zkscan3*-KO mESCs cultured in Serum/LIF or 2i/LIF over a ~3.5Mb region of chromosome 1. ChIP-seq signal from a reference ENCODE dataset (ENCFF001MYC) is shown for comparison. **(B)** Immunoblot for H3K9me3 levels relative to levels of unmodified Histone H3 in WT and *Zkscan3*-KO mESCs cultured in Serum/LIF or 2i/LIF. **(C)** Scatterplots showing normalised read counts under H3K9me3 peaks in WT and *Zkscan3*-KO mESCs cultured in Serum/LIF or 2i/LIF. Significantly differentially enriched peaks are shown in red. Pearson correlation between WT and *Zkscan3*-KO is indicated in the bottom right of each plot. Differential enrichment was assessed by DESeq2. **(D)** Genome browser tracks showing two significantly differentially enriched peaks in *Zkscan3* KO mESCs cultured in 2i/LIF. ZKSCAN3 ChIP-seq data is shown for reference.

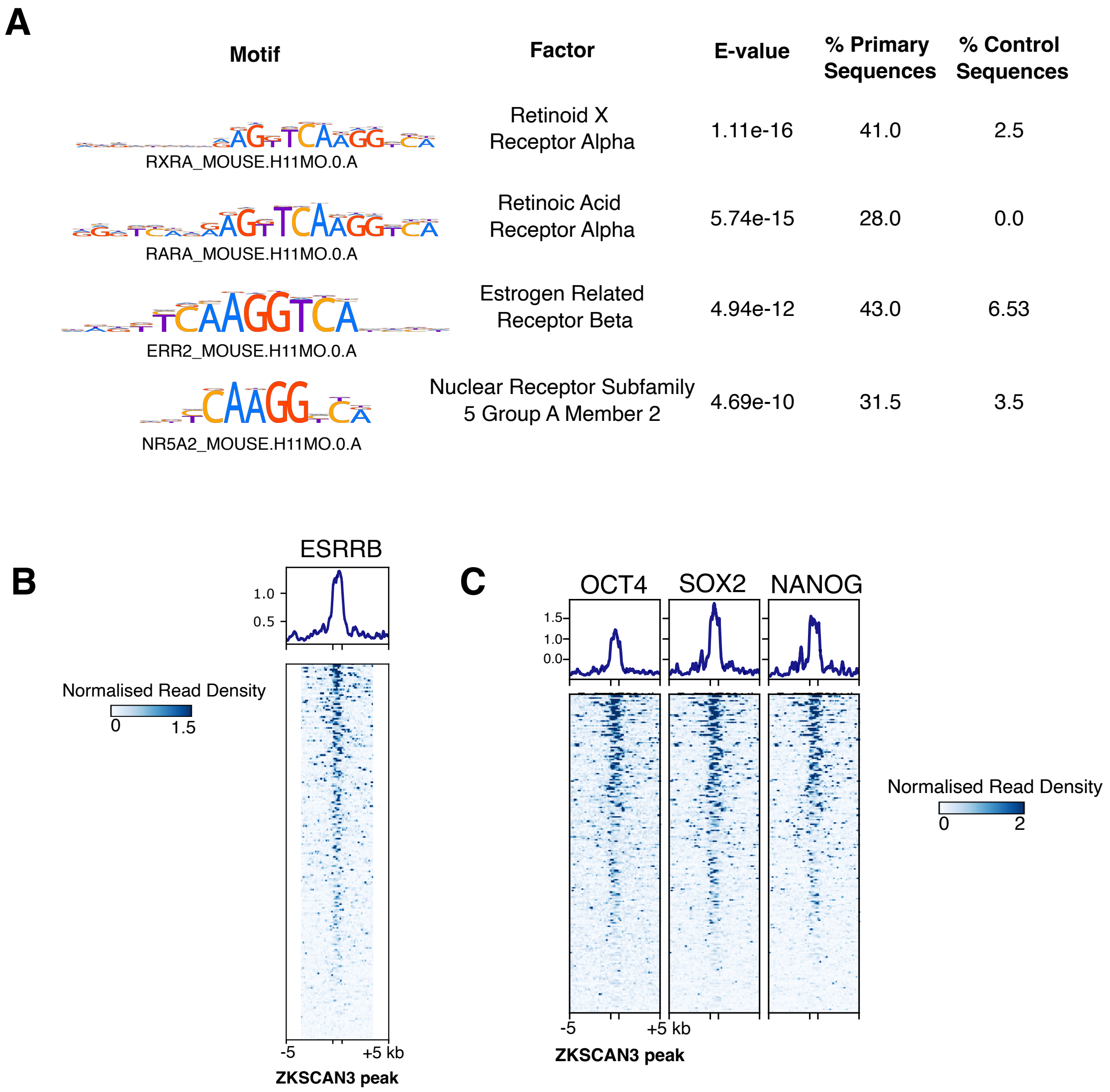


**Figure S6. ZKSCAN3 co-occupies many loci with pluripotency factors.**
**(A)** Top four motifs identified in ZKSCAN3 peak sequences. ZKSCAN3 peak sequences were scanned for known motifs occurring in the HOCOMOCO Mouse (v11 CORE) database. **(B)** Profile and heatmap of ESRRB ChIP-seq read density over ZKSCAN3 peaks. **(C)** Profile and heatmap of ChIP-seq read density over ZKSCAN3 peaks for pluripotency factors OCT4, SOX2 and NANOG.

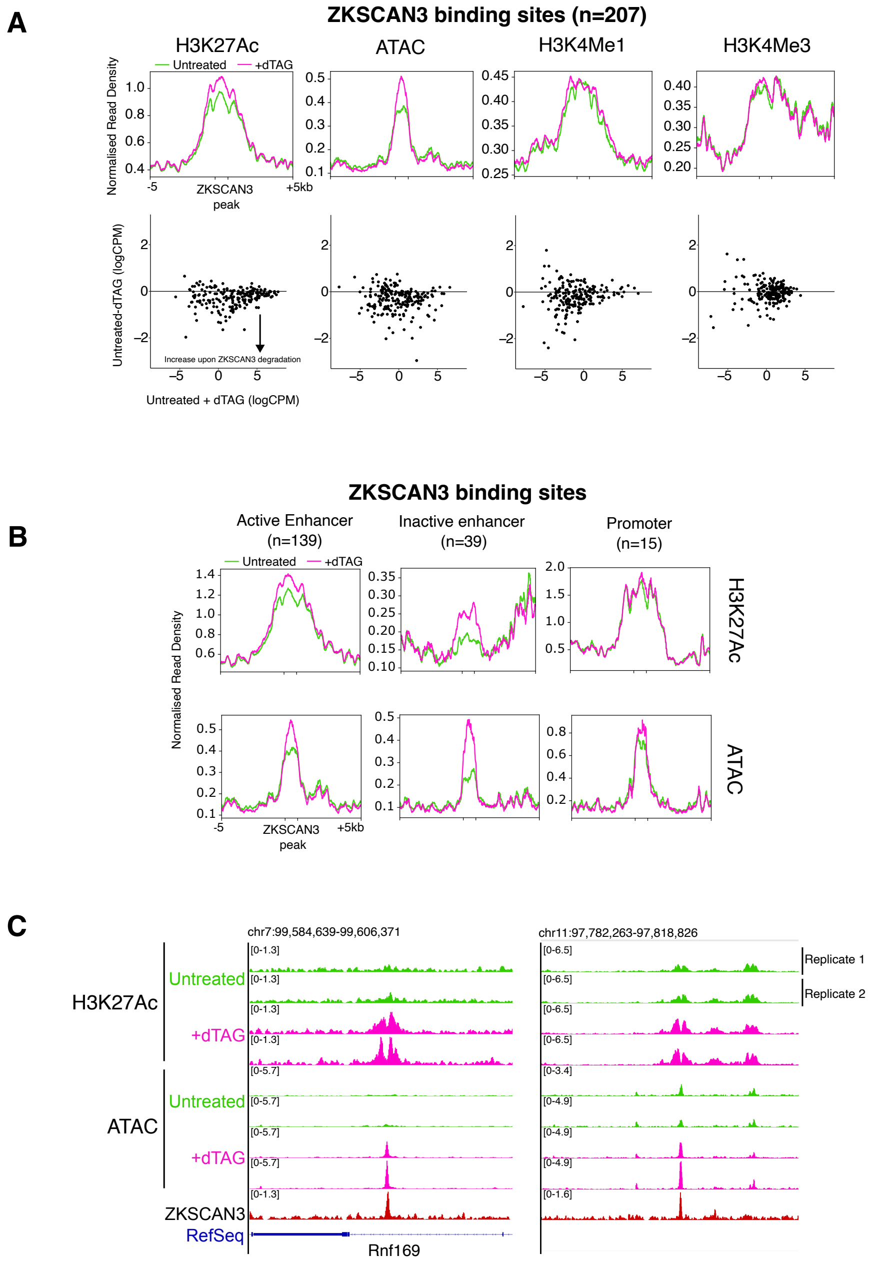


**Figure S7. ZKSCAN3 regulates H3K27 acetylation and chromatin accessibility**

**(A)** Metaplot profiles (above) and MA plots (below) showing differential enrichment of H3K27Ac, accessibility, H3K4me1 and H3K4me3 over ZKSCAN3 peaks, before and after ZKSCAN3 degradation with dTAG. **(B)** Metaplot profiles of H3K27ac (above) and ATAC (below) over ZKSCAN3 peaks subdivided into putative active enhancers, inactive enhancers and promoters based on the intersection of ZKSCAN3 peaks with H3K4me1/H3K27Ac, only H3K4me1 or H3K4me3, respectively, before and after ZKSCAN3 degradation with dTAG. **(C)** Genome browser track examples of an inactive (left) and an active (right) putative enhancer regulated by ZKSCAN3. For panels (A) and (B) each point on MA plots and the profile in metaplots show the average of two Zkscan3-Fkbpv-HA clonal line replicates. For metaplots, ZKSCAN3 peaks are scaled to 1000bp, and the region 5kb up and downstream is plotted for context.

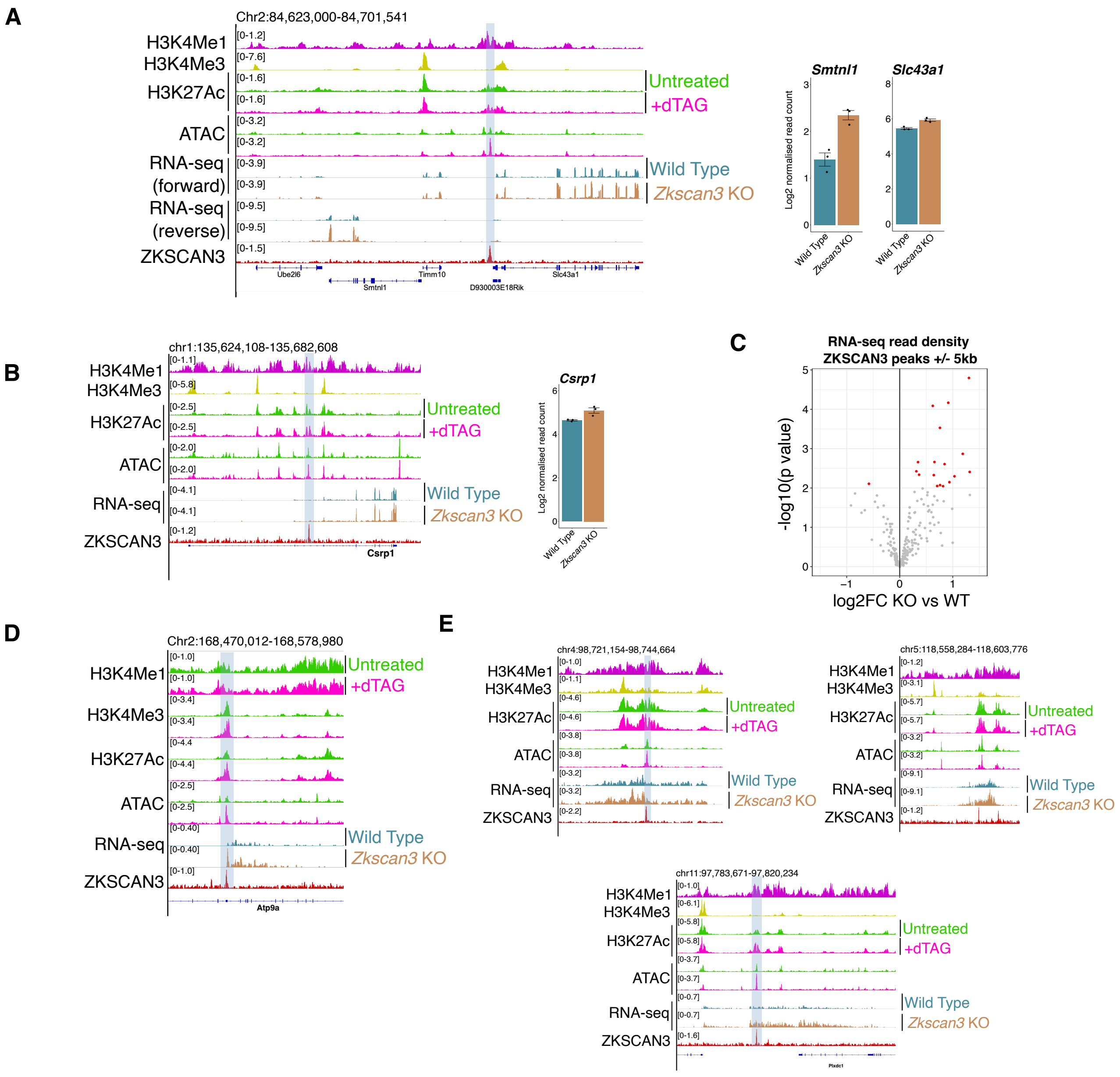


**Figure S8. ZKSCAN3 represses transcription in mESCs**

**(A)** Genome browser track example of a ZKSCAN3-bound putative enhancer, which gains accessibility upon ZKSCAN3 degradation and shows increased transcription of two proximal genes, *Slc43a1* and *Smtnl1* in *Zkscan3*-KO cells. **(B)** Genome browser track example of another ZKSCAN3-bound putative enhancer, which gains H3K27Ac and chromatin accessibility upon ZKSCAN3 degradation. The proximal gene *Csrp1* is upregulated in *Zkscan3*-KO mESCs. **(C)** Volcano plot showing the change in RNA-seq read density in *Zkscan3*-KO cells over and around ZKSCAN3 peaks (+/- 5kb). Significant hits ($p<0.05$) are shown in red. **(D)** Genome browser track example of a ZKSCAN3 bound promoter, which gains H3K4me3, H3K27ac and accessibility upon ZKSCAN3 degradation. An apparent lncRNA is up-regulated in *Zkscan3*-KO cells. **(E)** Genome browser track examples of un-annotated transcripts upregulated in *Zkscan3*-KO mESCs that intersect a ZKSCAN3 peak. For clarity, excepting panel (A), only RNA-seq reads in one direction are shown. Browser tracks show the average of 3 replicates (RNA-seq), 4 replicates (ZKSCAN3 ChIP-seq) or 2 replicates (ATAC-seq and other ChIP-seq) from individual clonal *Zkscan3-Fkbpv-HA* lines. In panels (A) and (B) the bar plots to the right of the browser tracks show the quantification of the indicated gene for all three RNA-seq replicates, in Log2 scale normalised to the read count.

A

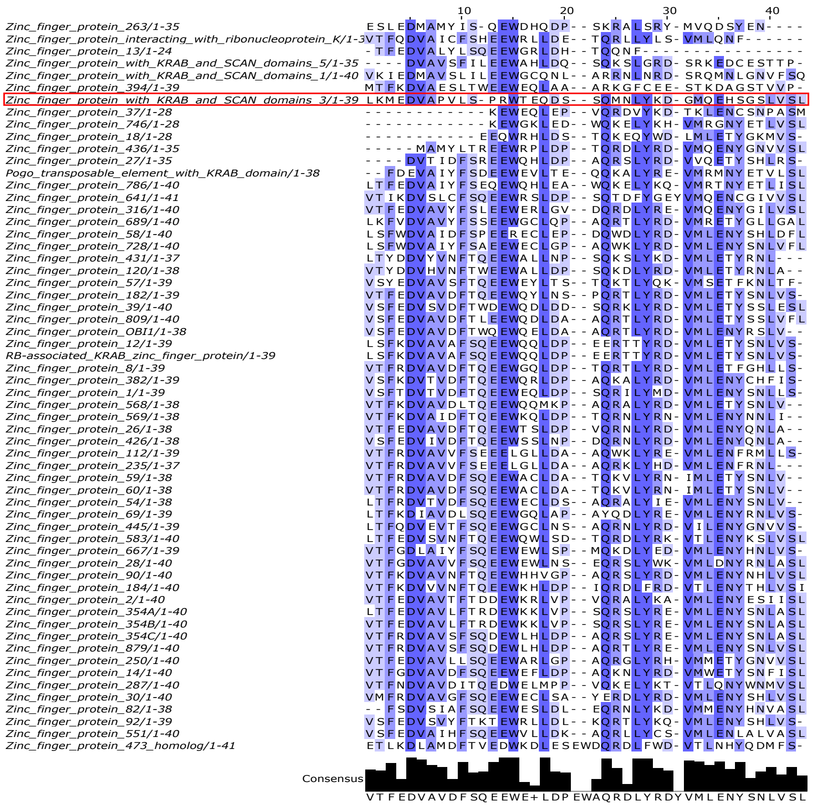


B

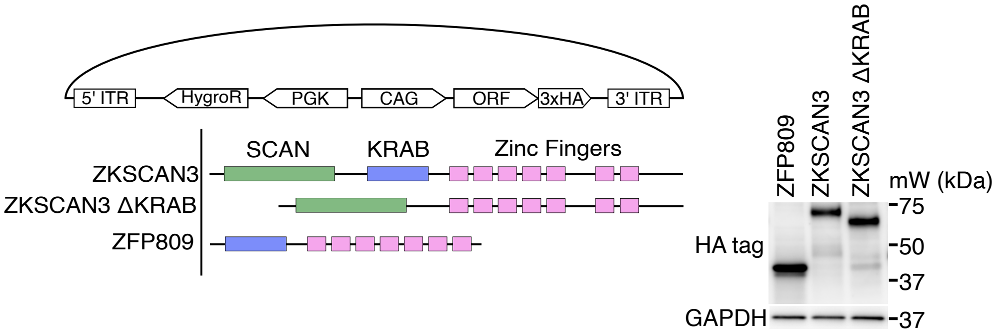


**Figure S9. ZKSCAN3 possesses a variant KRAB domain**

**(A)** Multiple sequence alignment of KRAB domain A-box sequences from InterPro. Sequences are ordered according to similarity as assessed by neighbour joining. The depth of shading indicates the level of conservation at each position. The consensus sequence is indicated at the bottom of the image. **(B)** Generation of ZKSCAN3, ZKSCAN3 ΔKRAB and ZFP809 over-expressing mESC lines. Open reading frames (ORFs) for the three sequences (bottom left) were cloned into a piggyBac expression cassette flanked by Inverted Terminal Repeats (ITRs) (top left) and transfected into mESCs along with piggyBac transposase to facilitate integration. Expression of all three constructs was confirmed by immunoblot (right). All proteins were HA-tagged to enable detection with the same antibody, and were generated at the same time from the same batch of mESCs.

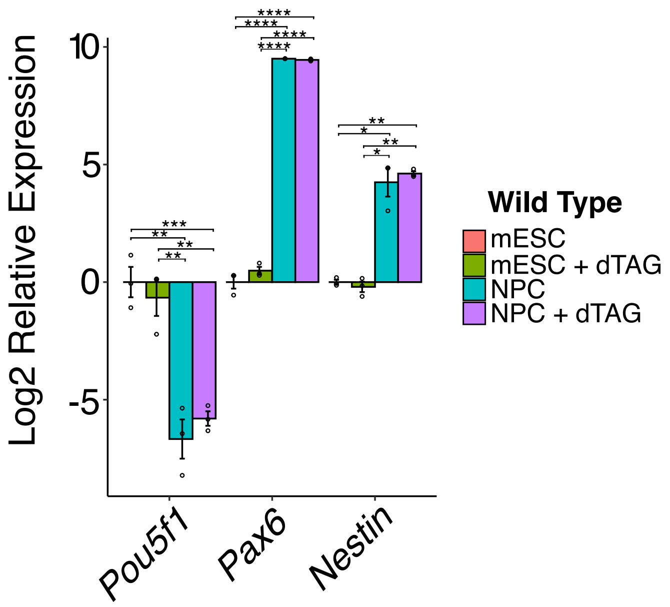


**Figure S10. dTAG does not affect neural differentiation of wild type mESCs**

RT-qPCR for the neural progenitor markers *Nestin* and *Pax6* and the pluripotency marker *Pou5f1* in WT mESCs differentiated towards the neural lineage, with and without the addition of dTAG. Expression is presented as the log2 relative expression to untreated mESCs. Expression is normalised to the geometric mean of *Rmr2* and *Gapdh*. Significance was assessed by one-way ANOVA followed by Tukey's post-hoc test. ****$p<1e-04$, ***$p<0.001$, **$p<0.01$, *$p< 0.05$.

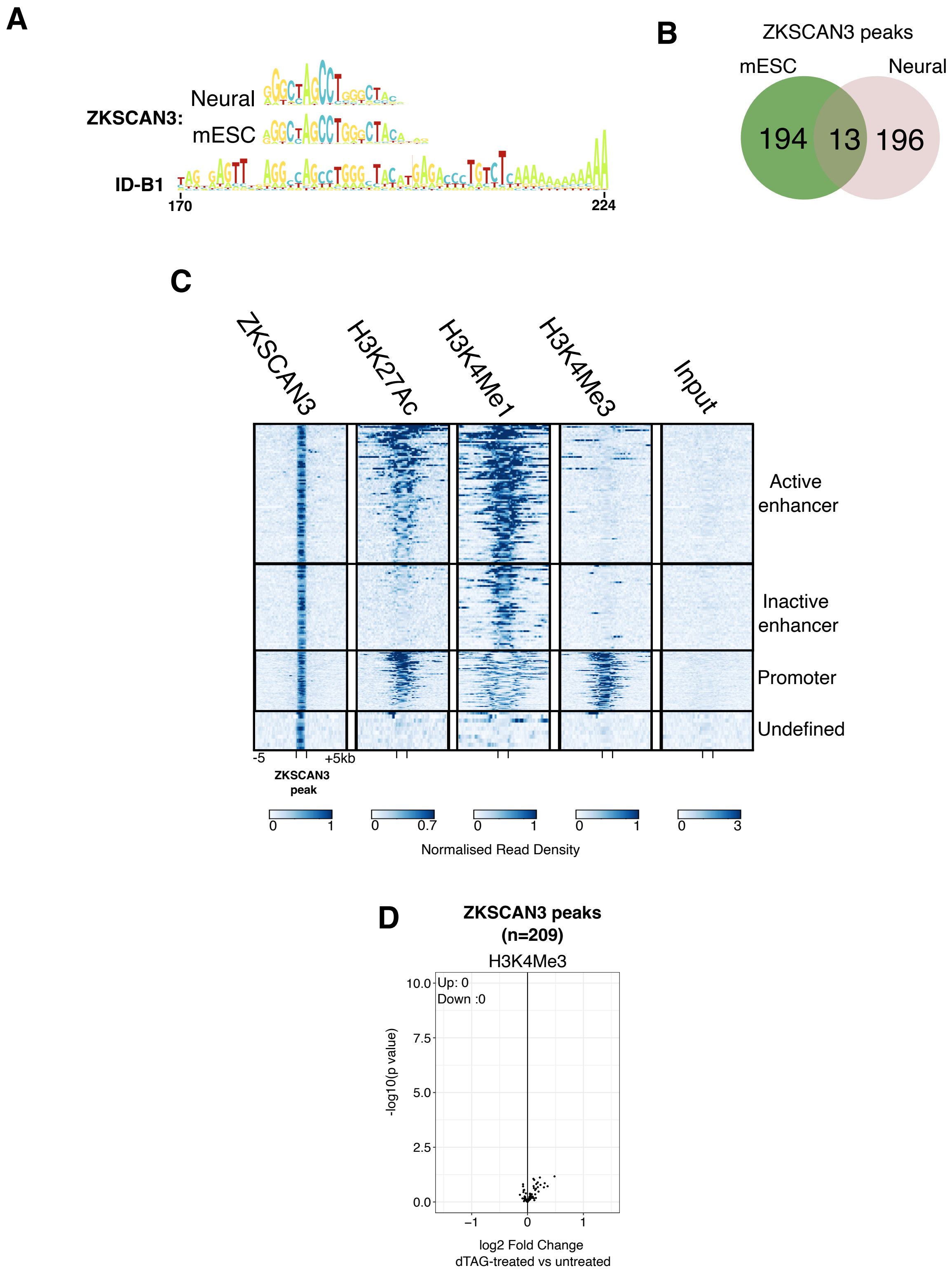


**Figure S11. ZKSCAN3 in the neural lineage**
**(A)** Motif identified in ZKSCAN3 ChIP-seq peaks in neural cells. The motif recovered from ChIP-seq in mESCs, and the 3' end of the ID_B1 SINE element are presented for comparison. **(B)** Venn diagram showing the number of common ZKSCAN3 binding loci in mESCs and neural cells. **(C)** Heatmap showing normalised read density of ZKSCAN3, H3K27ac , H3K4me1 and H3K4me3 ChIP-seq over ZKSCAN3 peaks. ZKSCAN3 peaks are scaled to 1000bp and heatmaps show 5kb up and downstream. **(D)** Volcano plot showing the change in enrichment of H3K4me3 at ZKSCAN3 binding sites in the neural lineage. No significant changes ($p<0.05$) were detected. Each point represents the average of two replicates.

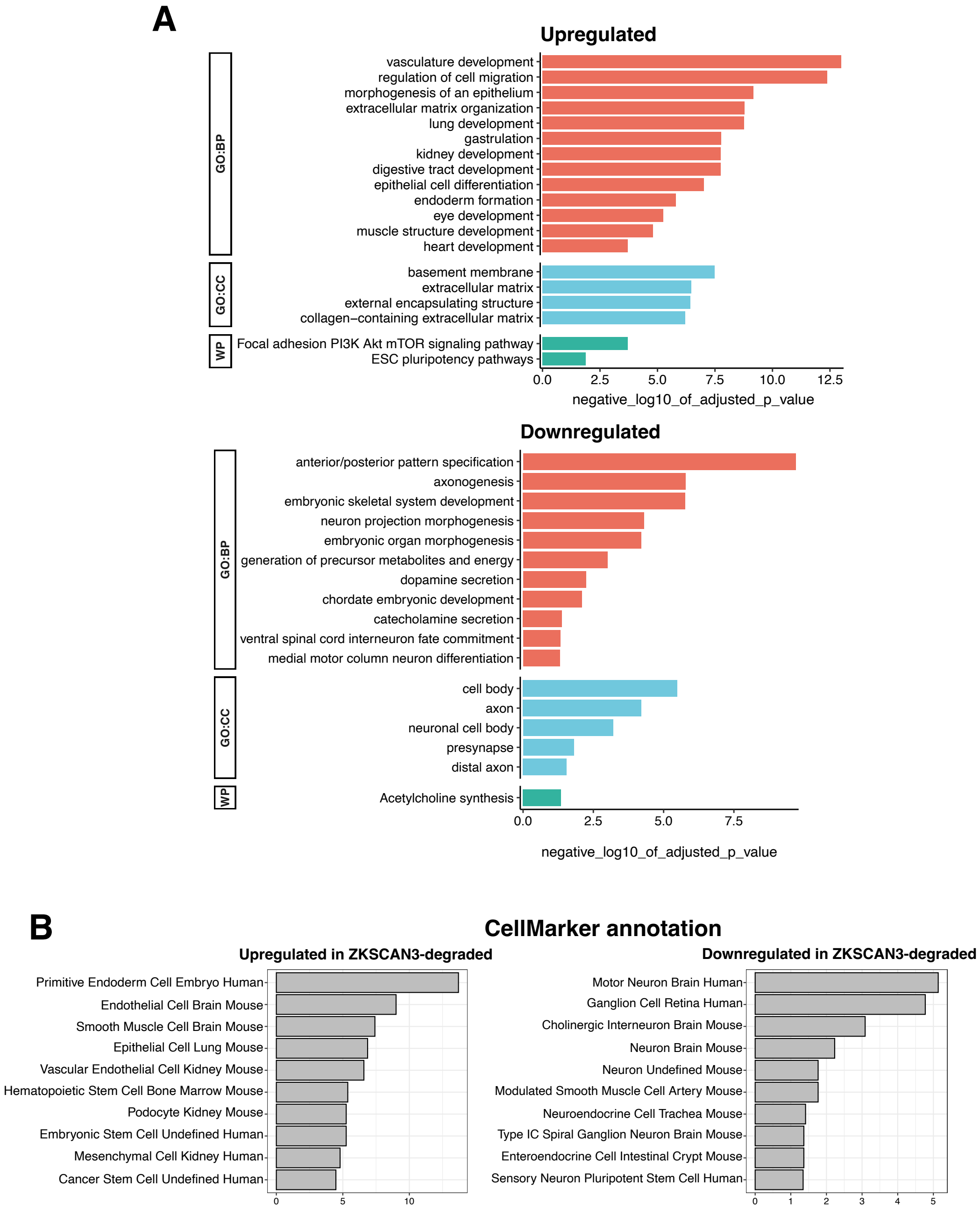


**Figure S12. Gene expression changes in ZKSCAN3-degraded neural cells**

**(A)** Gene Ontology (GO) analysis of genes upregulated (above) and downregulated (below) in ZKSCAN3-degraded neural cells. For clarity, and to remove highly similar terms, only select significant terms are shown; the full GO results are available in Supplementary Table 4. **(B)** CellMarker annotation of genes up- and down-regulated in the ZKSCAN3-degraded neural cells. The top 10 most significant matches to CellMarker annotation are presented. BP = Biological Process; CC = Cellular Component; WP = WikiPathways

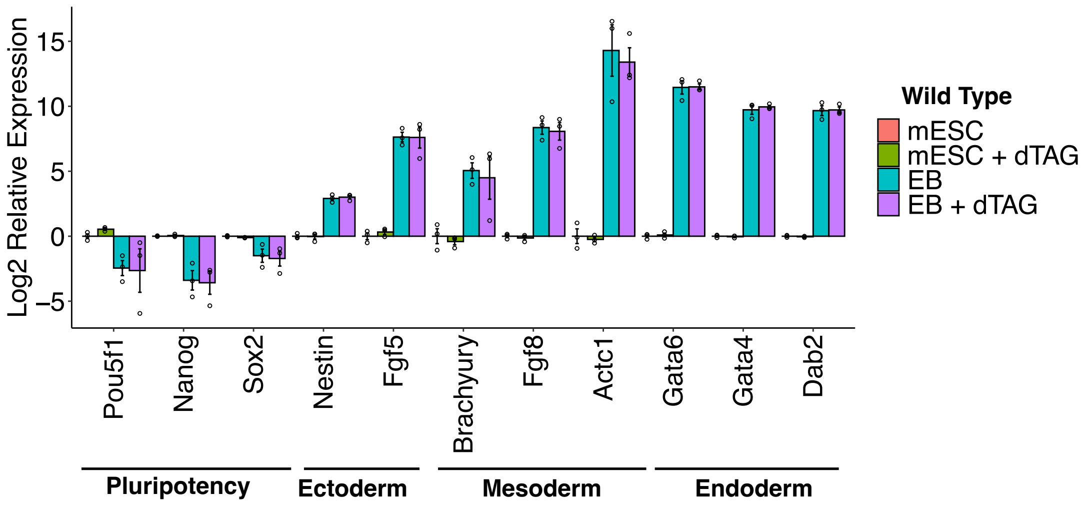


**Figure 13. dTAG alone does not affect differentitation of mESCs to embryoid bodies**

RT-qPCR of key pluripotency (*Pou5f1*, *Nanog*, *Sox2*), ectodermal (*Nestin*, *Fgf5*), mesodermal (*Brachyury*, *Fgf8*, *Actc1*) and endodermal (*Gata6*, *Gata4*, *Dab2*) markers following embryoid body formation from WT mESCs in the presence or absence of dTAG. Expression is calulated relative to untreated mESCs. Data are normalised to the expresion level of *Rmr2*. Error bars show SEM.

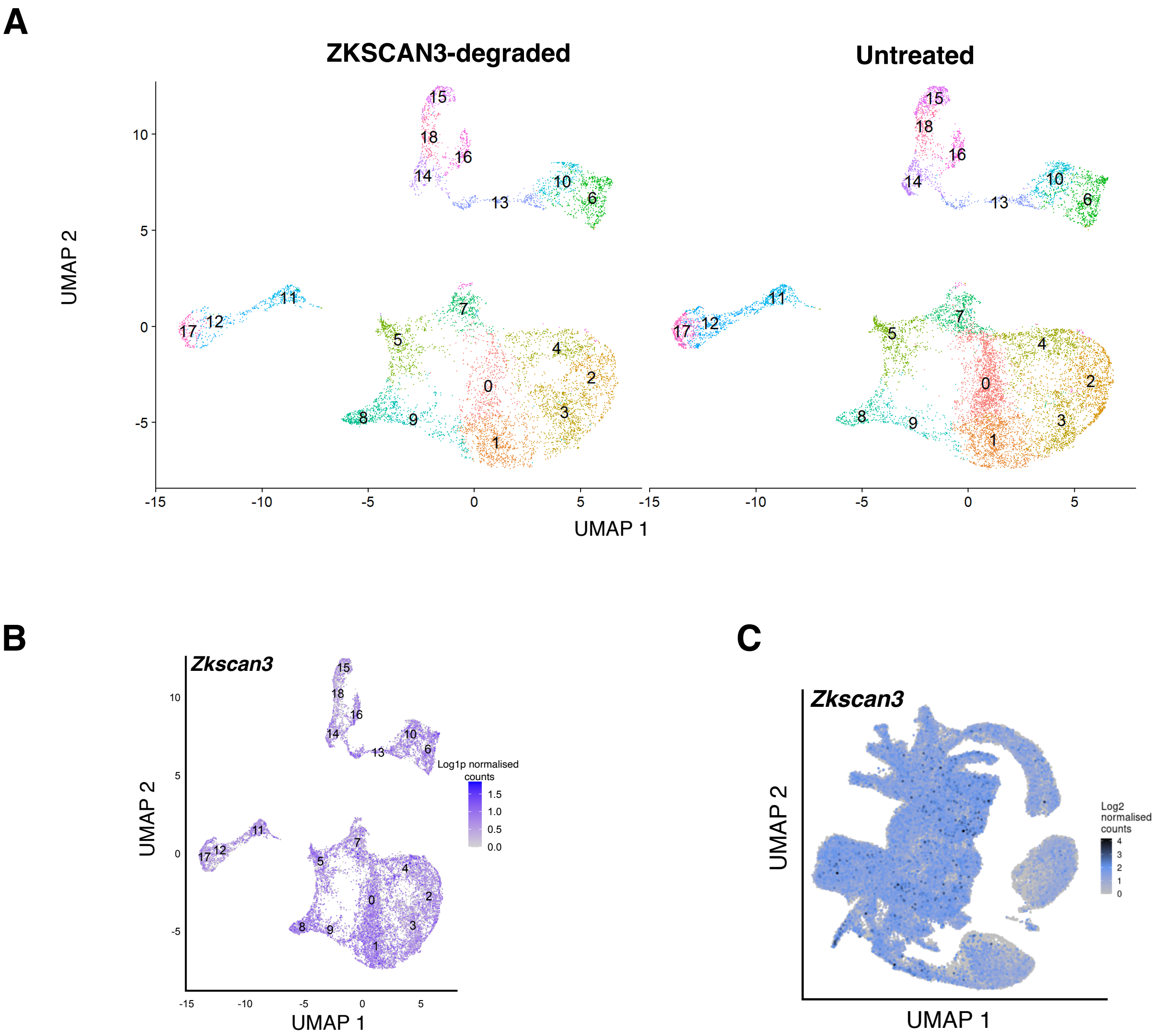


**Figure S14. scRNA-seq of embryoid bodies**

**(A)** UMAP projection of cells in untreated and ZKSCAN3-degraded Embryoid Bodies (EBs). Cells are clustered and numbered by Seurat cluster. **(B)** UMAP projection of *Zkscan3* expression in untreated EBs and **(C)** in the mouse embryo, from The Mouse Gastrulation Atlas.

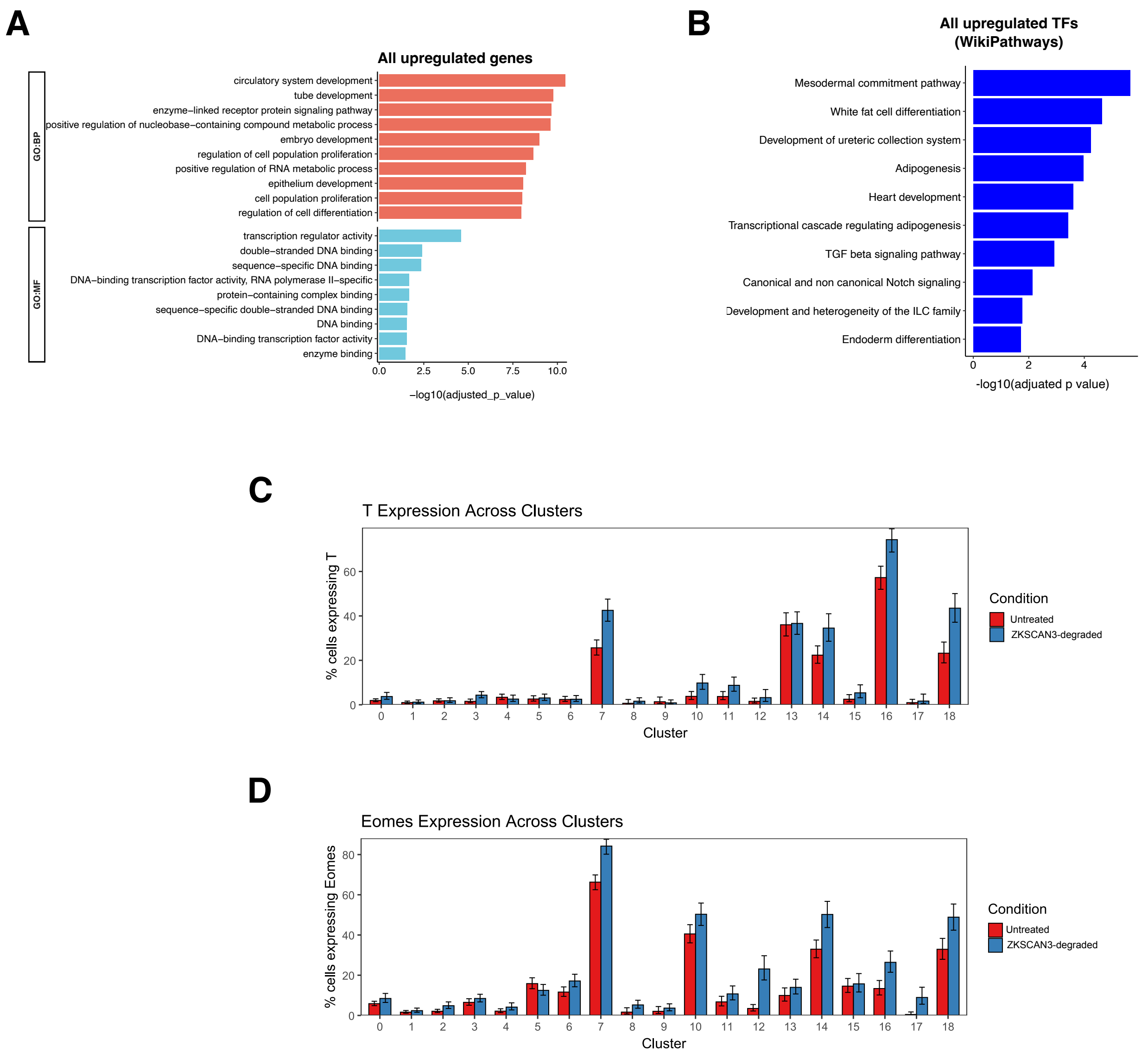


**Figure S15. Upregulated genes in ZKSCAN3-degraded EBs are enriched for mesoendodermal TFs**

**(A)** GO analysis of all upregulated genes in ZKSCAN3-degraded EBs. **(B)** GO analysis presented against the WikiPathways database for all upregulated transcription factors in ZKSCAN3-degraded EBs. **(C)** Bargraph showing expression of *T* (*brachyury*) and **(D)** *Eomes* expression across clusters in untreated and ZKSCAN3-degraded EBs. Error bars show 95% confidence intervals. For GO analysis, only GO terms with <1000 members are shown. MF = Molecular Function; BP = Biological Process

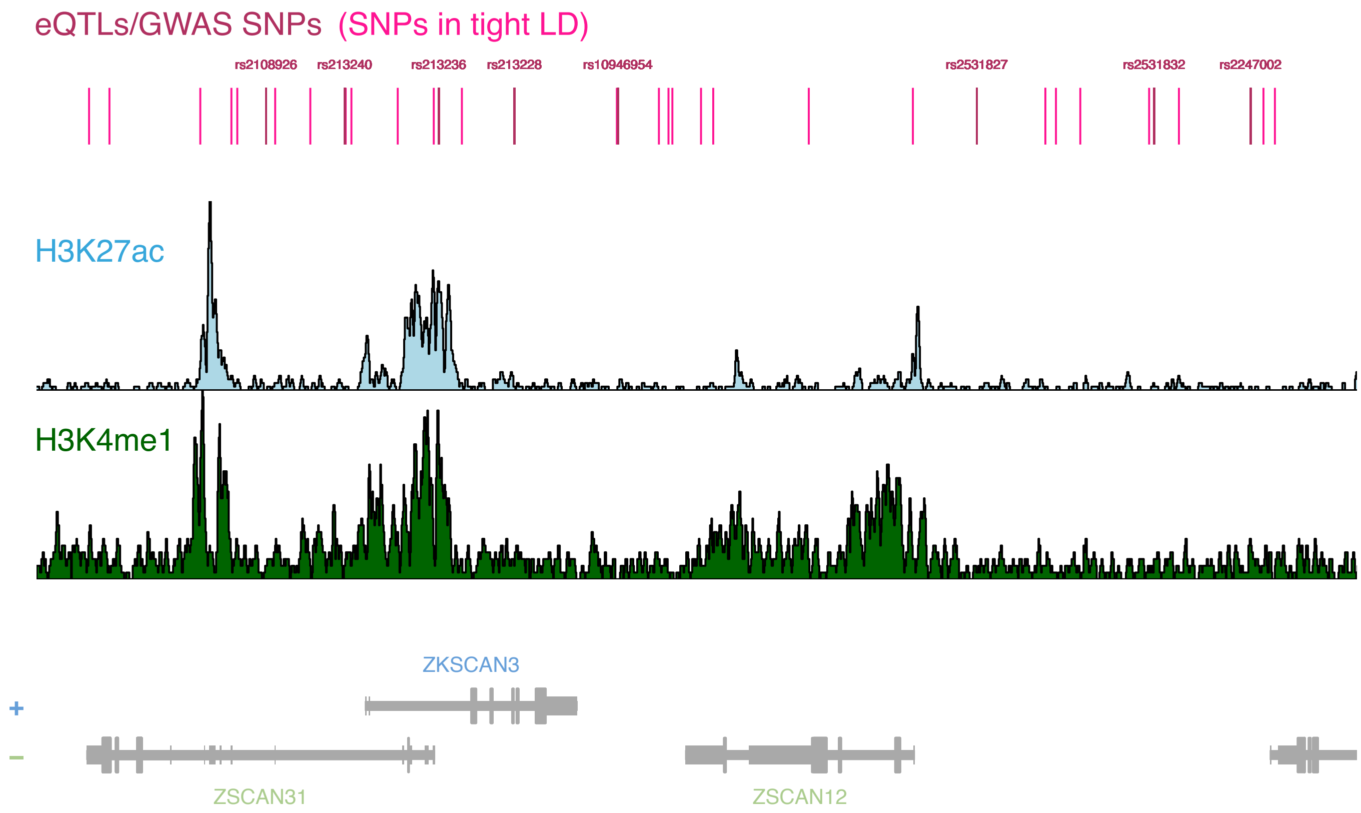


**Figure S16. SNPs assosciated with SZ risk and gray matter volume changes are eQTLs for *ZKSCAN3* in human brain**

Schematic showing the location of 8 SNPs identified in Chen et al[31] that are associated with grey matter volume changes and are eQTLs for *ZKSCAN3*. SNPs are shown in red, and SNPs in tight linkage disequilibrium with these SNPs ($R^2$>0.9) are shown in pink. Histone modification ChIP-seq data for the human mid-frontal lobe region are presented below to highlight putative enhancer regions.
